# Origins of proprioceptor feature selectivity and topographic maps in the *Drosophila* leg

**DOI:** 10.1101/2022.08.08.503192

**Authors:** Akira Mamiya, Anne Sustar, Igor Siwanowicz, Yanyan Qi, Tzu-Chiao Lu, Pralaksha Gurung, Chenghao Chen, Jasper S. Phelps, Aaron T. Kuan, Alexandra Pacureanu, Wei-Chung Allen Lee, Hongjie Li, Natasha Mhatre, John C. Tuthill

## Abstract

Our ability to sense and move our bodies relies on proprioceptors, sensory neurons that detect mechanical forces within the body. Proprioceptors are diverse: different subtypes detect different features of joint kinematics, such as position, directional movement, and vibration. However, because they are located within complex and dynamic peripheral tissues, the underlying mechanisms of proprioceptor feature selectivity remain poorly understood. Here, we investigate molecular and biomechanical contributions to proprioceptor diversity in the *Drosophila* leg. Using single-nucleus RNA sequencing, we found that different proprioceptor subtypes express similar complements of mechanosensory and other ion channels. However, anatomical reconstruction of the proprioceptive organ and connected tendons revealed major biomechanical differences between proprioceptor subtypes. We constructed a computational model of the proprioceptors and tendons, which identified a putative biomechanical mechanism for joint angle selectivity. The model also predicted the existence of a goniotopic map of joint angle among position-tuned proprioceptors, which we confirmed using calcium imaging. Our findings suggest that biomechanical specialization is a key determinant of proprioceptor feature selectivity in *Drosophila*. More broadly, our discovery of proprioceptive maps in the fly leg reveals common organizational principles between proprioception and other topographically organized sensory systems.

## Introduction

Flexible motor control of the arms and legs requires sensory feedback from proprioceptive sensory neurons (i.e., proprioceptors). Both invertebrate and vertebrate animals possess multiple subtypes of proprioceptors that detect unique aspects of limb position and movement (Tuthill and Azim, 2018). For example, in mammalian muscle spindles, Group Ia and II afferents encode muscle length and velocity (Macefield and Knellwolf, 2018). A functionally analogous structure in the insect leg, the femoral chordotonal organ (FeCO), monitors the kinematics of the femur-tibia joint (Field and Matheson, 1998; Krishnan and Sane, 2015). The FeCO of the fruit fly, *Drosophila melanogaster*, contains three subtypes of mechanosensory neurons with distinct stimulus feature selectivity and axonal projections (Mamiya et al., 2018): claw neurons encode tibia position (flexion or extension), hook neurons encode directional movement (flexion or extension), and club neurons encode bidirectional movement and low amplitude, high-frequency vibration (**Fig. 1**). Claw and hook neurons likely contribute to feedback control of leg movements, such as walking and grooming, while club neurons may be used to monitor vibrations in the external environment, such as detecting conspecific wingbeats during social interactions (Agrawal et al., 2020; Chen et al., 2021; McKelvey et al., 2021).

**Fig. 1.**
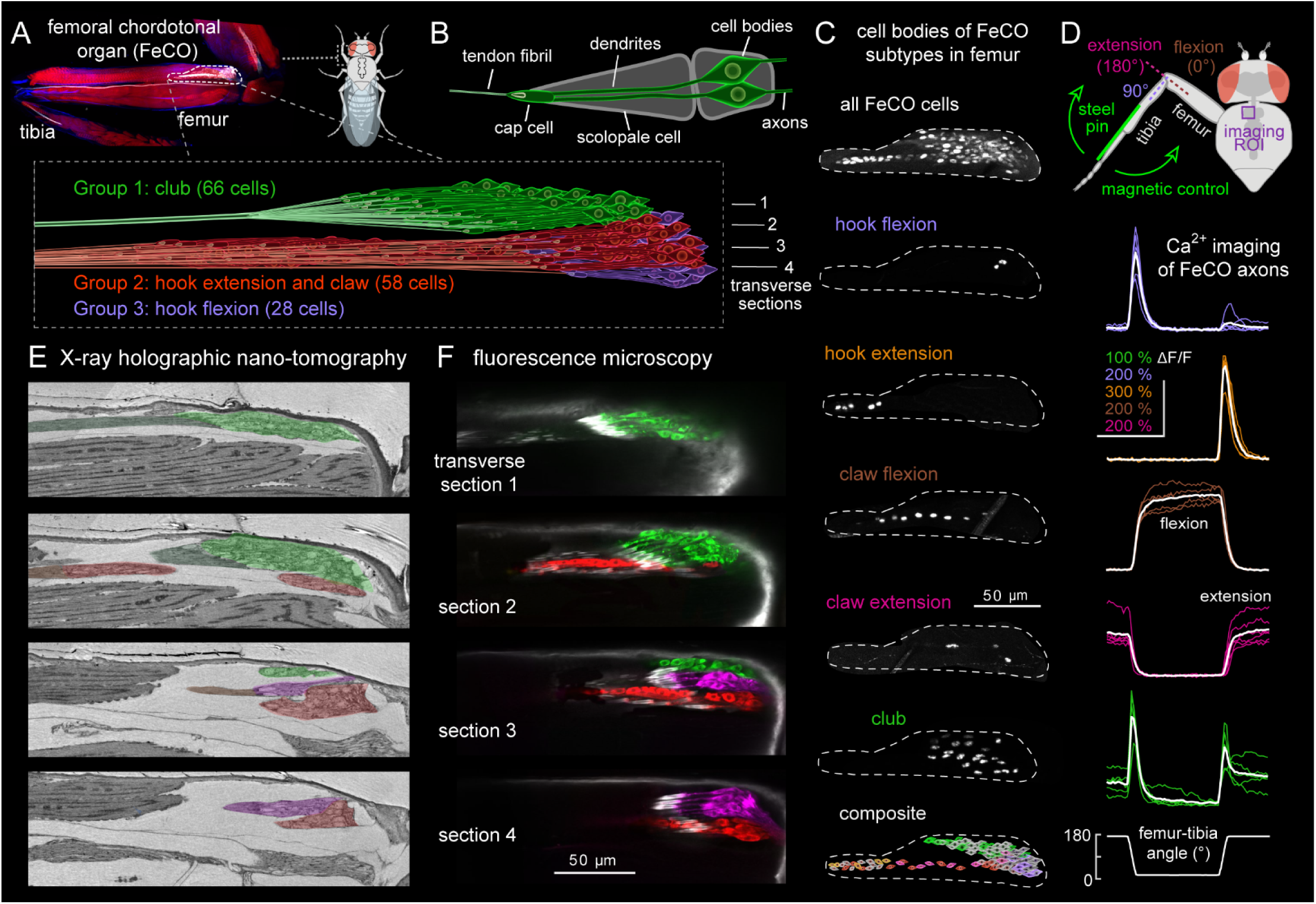
Functional subtypes of FeCO proprioceptors are spatially clustered within three discrete compartments in the *Drosophila* leg. **(A)** Top: peripheral anatomy of the femoral chordotonal organ (FeCO, labeled with GFP driven by *iav-Gal4* (white); red is a phalloidin stain; blue is cuticle autofluorescence. Bottom: schematic of FeCO organization, based on X-ray reconstruction. **(B)** Dendrites of each pair of FeCO cells are surrounded by a scolopale cell that connect to a cap cell, which in turn connects to a tendon. (**C**) Split-Gal4 lines driving RFP label subtypes of FeCO cell bodies in specific locations in the femur. Bottom: a composite schematic showing the relative locations of cell bodies for each FeCO subtype. **(D)** Two-photon calcium imaging from axons of each FeCO subtype during controlled movements of the femur-tibia joint. Thin traces are from individual flies (n = 5-7) and the thick white line is the response average. **(E)** Reconstruction from an X-ray microscopy dataset of the fly femur reveals FeCO organization. Each image corresponds to a transverse section indicated in the schematic above. FeCO compartments and tendons are indicated by color shading. **(F)** The same image planes as **C** but visualized with confocal imaging (GFP driven by *iav-Gal4*; pseudo-colored to indicate compartment). Chordotonal cap cells are labeled with an antibody against phalloidin (white).

What mechanisms determine the feature selectivity of different proprioceptor subtypes? One possibility is that differences in intrinsic cellular properties, such as expression of specific ion channels, allow different sensory neuron subtypes to sense distinct mechanical features. In mammals, muscle spindle sensory neurons sense mechanical forces using the Piezo2 channel (Nagel and Chesler, 2022), with possible contributions from DEG/ENaC channels (Bewick and Banks, 2021). Single-cell RNA sequencing from muscle spindle and golgi tendon sensory neurons exhibit diversity in the expression of voltage gated potassium channels, but not mechanosensory ion channels (Oliver et al., 2021). Chordotonal neurons in *Drosophila* express the mechanosensory channels Piezo, NompC, Iav, Nan, and Painless (Hehlert et al., 2021), although the distribution of these and other ion channels across proprioceptor subtypes is not known.

An alternative mechanism for proprioceptor feature selectivity is biomechanical differences. For example, specialized attachment structures could decompose and transmit distinct forces to different proprioceptor subtypes. Indeed, the name *chordotonal* was originally coined by Vitus Graber in the 19^th^ century, based on his observation of cells that resemble a bundle of chords under tension (Graber, 1881). However, it remains unknown how the *chords* (i.e., tendons) contribute to tuning of mechanosensory neurons. The experimental inaccessibility of proprioceptive sensory organs, which are typically embedded deep within the body, have prevented investigation of their mechanical operation *in vivo*.

Here, we develop and apply new experimental methods to investigate the relative contributions of molecular and biomechanical mechanisms to proprioceptor feature selectivity and topographic encoding in the *Drosophila* leg. Using single-nucleus RNA-sequencing, we found that proprioceptor subtypes are transcriptionally distinct, but lack differential expression of mechanosensory ion channels. We combined anatomical reconstruction and biomechanical modeling to reveal that different proprioceptor subtypes are positioned to receive different mechanical signals from the same joint. We also discovered that the cell bodies of position-tuned proprioceptors contain a *goniotopic* map of joint angle, whereas vibration-tuned proprioceptors contain a *tonotopic* map of tibia vibration frequency. Our results motivate the investigation of peripheral biomechanics and topographic maps in proprioceptors of other limbed animals.

## Results

### The *Drosophila* FeCO is organized into three anatomical compartments

We reconstructed the peripheral ultrastructure of the FeCO using an X-ray holographic nano-tomography dataset of the *Drosophila* front leg (Kuan et al., 2020). We identified 152 total cell bodies in the FeCO, which are organized into three separate compartments called scoloparia (Field and Matheson, 1998) (**Fig. 1, Movies S1, S2)**. We constructed split-Gal4 genetic driver lines that specifically label club, claw, and hook neurons (**Fig. 1C-D, S1, S2, Movie S3**) and compared fluorescence and X-ray images to map the peripheral location of each FeCO subtype within the femur (**Fig. 1C, E-F**, **S1B-D**). We found that the largest and most dorsal compartment (Group 1) contains the club neurons, which encode tibia vibration and cyclic movement. Claw neurons, which encode the static angle of the femur-tibia joint, are distributed in a linear array along the long axis of the femur (Group 2). A smaller and more lateral compartment (Group 3) contains the hook flexion neurons, which respond transiently when the tibia flexes, but not during extension. The hook extension neurons, which respond transiently during tibia extension but not flexion, are located distal to the claw neurons in Group 2. Overall, our reconstruction revealed that FeCO neurons are organized into three anatomical compartments within the femur, suggesting that they may experience different forces as a result of tibia joint movements.

### FeCO subtypes are transcriptionally distinct but express overlapping subsets of mechanosensory ion channels

We next used single-cell RNA sequencing to ask if differences in gene expression among proprioceptor subtypes could explain the differences in their mechanical feature selectivity (**Fig. 2**). We genetically labeled FeCO neurons with GFP and sequenced nuclei that showed green fluorescence above a specific threshold (**Fig. 2A**). Clustering of gene expression revealed an enrichment of putative FeCO nuclei, but also inclusion of other cell-types, presumably due to low-levels of auto-fluorescence or some droplets consisting of multiple nuclei during sorting (**Fig. 2B, C**). We were able to distinguish neuronal and non-neuronal clusters based on the expression levels of known cell markers (**Fig. 2C, Table S1;** (Li et al., 2022)). For the three putative clusters of FeCO neurons (**Fig. 2D)**, we used gene-specific Gal4 lines to validate the expression of the top candidate genes differentially expressed in each cluster (**Fig. S3A**). This revealed that the three clusters correspond to club, claw, and hook neurons. We were unable to resolve distinct clusters for extension and flexion-tuned hook and claw neurons, either because they are transcriptionally similar or because their differences are below the resolution of our analysis.

**Fig. 2.**
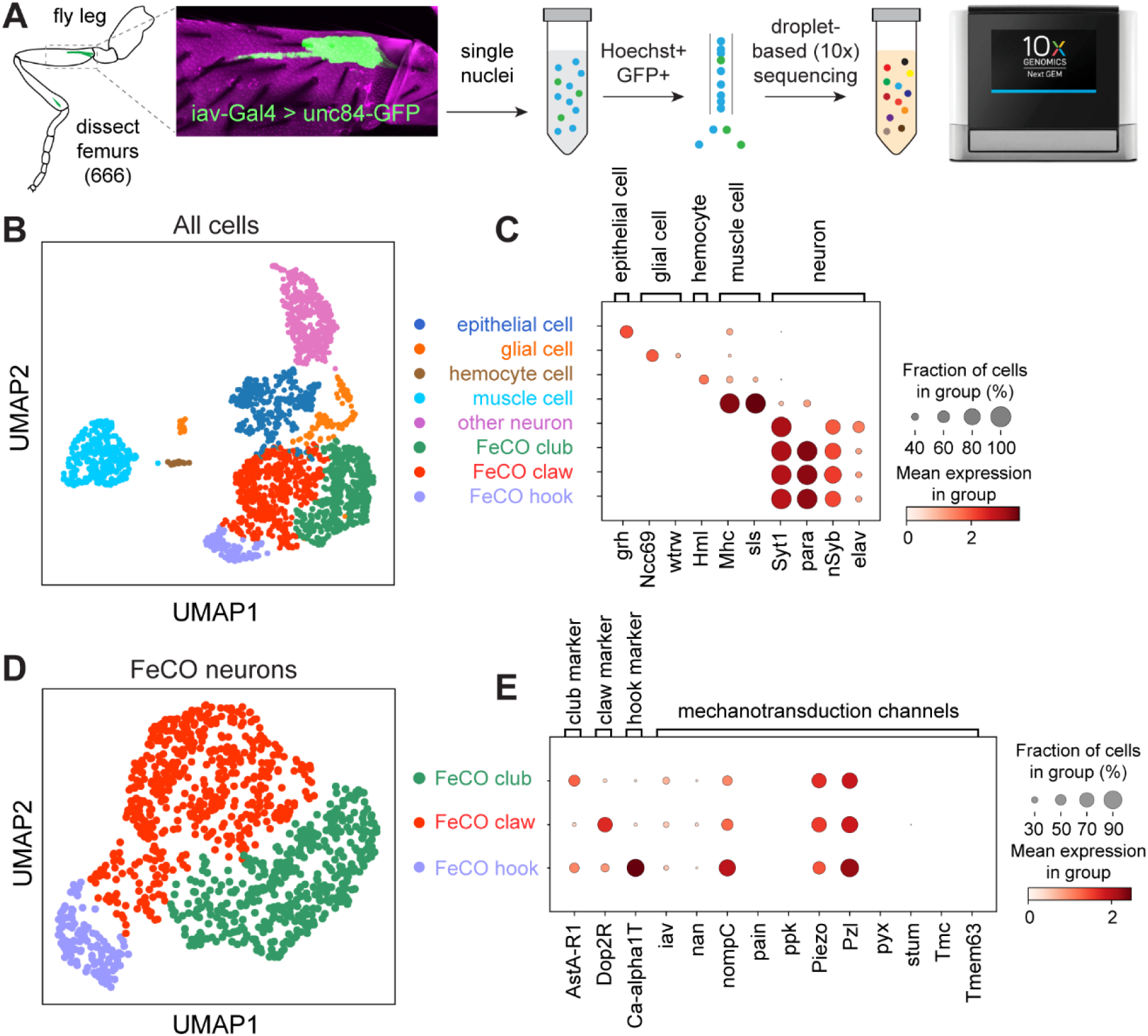
Single-nucleus RNA-sequencing reveals overlapping expression of multiple mechanotransduction channels among FeCO subtypes. (**A**) Femurs from 666 fly legs expressing GFP were dissected and nuclei were extracted and collected using FACS for droplet-based (10x) sequencing. (**B**) UMAP visualization of eight clusters with annotated cell types. See methods for clustering details. (**C**) Dot plot showing expression levels of cell-type and neuronal markers in each of the eight clusters. (**D**) UMAP of the three FeCO subtypes from **B**. (**E**) **Left:** Dot plot showing expression levels of cell-type candidate genes for FeCO club (AstaAR1), claw (Dop2R), and hook (Ca-alpha1T). **Right:** Mechanotransduction channel expression for each of the three FeCO clusters. In dot plots, color intensity represents mean level of gene expression in a cluster relative to the level in other clusters, and size of dots represents the percent of nuclei in which gene expression was detected.

We next asked whether each cluster expresses different mechanosensory ion channels, which could provide a mechanism for feature selectivity among FeCO subtypes. However, we found that known mechanosensory channel RNAs were expressed similarly on average across the three FeCO clusters (**Fig. 2E, S3B**). We also examined the expression profiles of voltage gated sodium and potassium channels and again found no differences in channel expression (**Fig. 2C, S4**). These results suggest that differential expression of known mechano-sensory and voltage-gated ion channels is unlikely to account for feature selectivity of proprioceptor subtypes. Because our analysis could not detect expression of different protein isoforms, we cannot rule out a role for alternative splicing of channel genes, as has been shown for the tonotopic map of hair cells in chicken cochlea (Fettiplace and Fuchs, 1999).

**Fig. 4.**
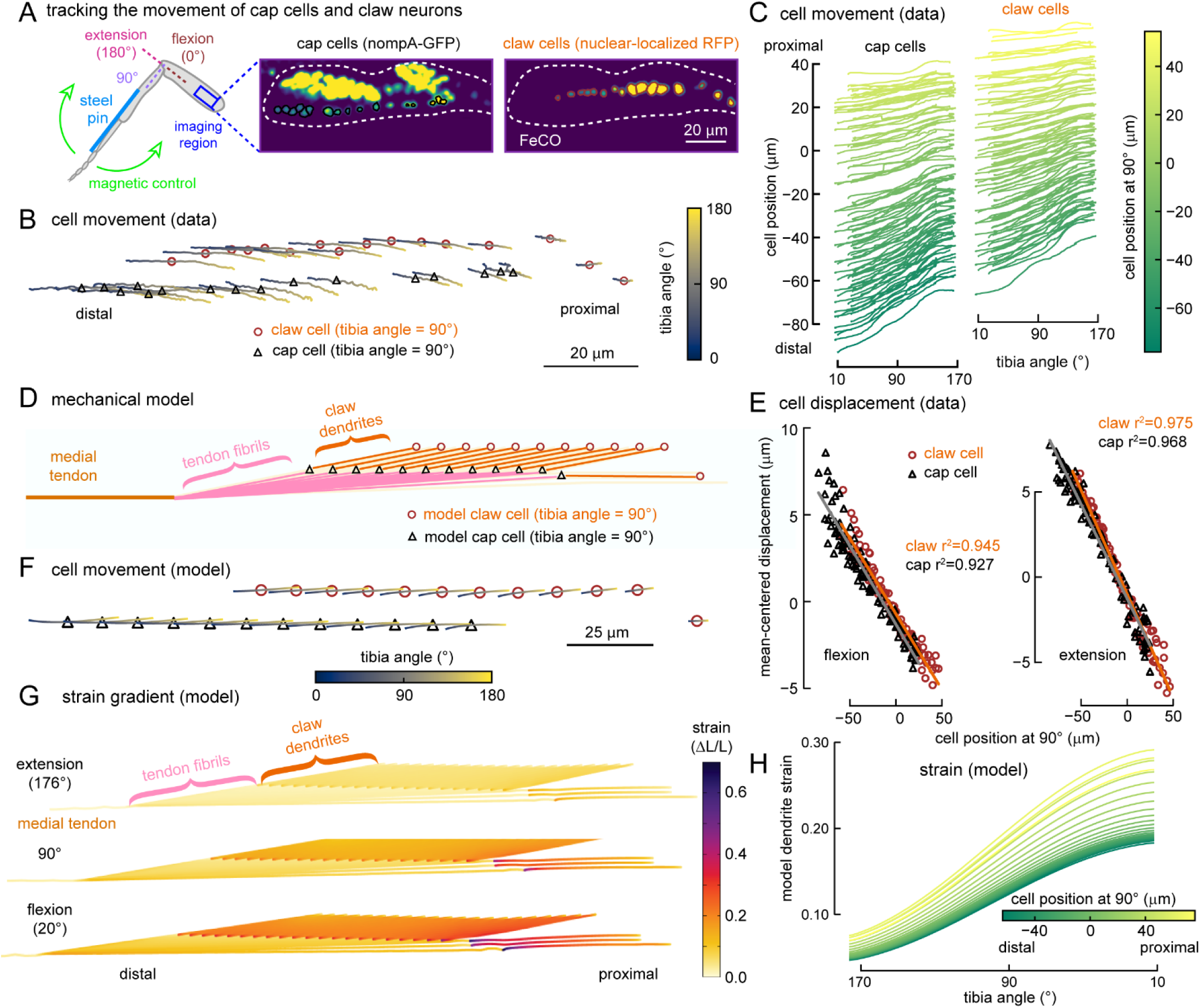
A spatial gradient of cell movement and mechanical strain among position-tuned claw neurons. **(A)** We imaged RFP labeled claw cell nuclei and GFP labeled cap cells (each colored circle indicates the tracked cell) in the femur using transcuticular twophoton microscopy, while swinging the tibia from extension to flexion with a magnetic control system. (**B**) Example traces showing the position of claw cells (brown circles) and cap cells (grey triangles) during full tibia flexion. The color of each trace indicates tibia angle. (**C**) The position of cap cells (left) and claw cells (right) along the distal-proximal axis of the femur as the tibia moved from full extension to full flexion (n = 6 flies). We mean-centered the cell position within each fly by subtracting the average claw cell position when tibia is at 90° for each fly. **(D)** A finite element model of claw and cap cells. Model claw cells connect to the cap cells via their dendrites. The cap cells in turn connects to the medial tendon via tendon fibrils that fan out from the end of the medial tendon. Only 1/2 of the cells in the model are shown for display purposes. **(E)** Movement of claw and cap cells during tibia flexion (left) and extension (right) vs the cell’s position along the distal to proximal axis of the femur (n = 6 flies). We mean-centered the cell displacement within each fly by subtracting the average claw displacement for each fly. (**F**) Same as **C**, but for the movement of cells in the model. **(G)** A map of strain in the dendrites of model claw cells at different tibia angles. (**H**) The strain (1st principal invariant strain) in different claw dendrites plotted against the tibia angle. The dendrites stretched and strain increased as tibia flexed.The strain was always larger in proximal cells and the difference increased as the tibia was flexed. The color of each line represents the cell position along the distal-proximal axis of the femur when tibia is at 90°.

### FeCO proprioceptors connect to the tibia through different tendons

We next asked whether there exist biomechanical differences between proprioceptor subtypes. To understand how FeCO neurons are mechanically coupled to the tibia, we used the X-ray microscopy dataset to reconstruct their distal attachment structures. The ciliated dendrites of each pair of FeCO neurons are ensheathed in a scolopale cell that attaches to a sensory tendon via an actin-rich cap cell (**Fig. 1B**). Tracing the termination point of each cap cell allowed us to determine the tendon attachment of each FeCO neuron in the X-ray and fluorescence images (**Fig. 1E-F, Movie S2**). We found that the three different FeCO compartments attach to different tendons (**Fig. 3A**). The tendons connected to Groups 2 and 3 (claw and hook) merge and fuse ∼100 μm distal to the FeCO, while the tendon connected to Group 1 (club) remained separate. The two tendons, which we refer to as medial and lateral, then converge upon a tooth-shaped structure in the distal femur (**Fig. 3A-C, Movie S4**). A similar structure was previously described in the beetle leg, where it was named the *arculum* (Frantsevich et al., 2019). In *Drosophila*, we found that the medial tendon attaches to the arculum’s medial root, while the lateral tendon attaches to the arculum’s lateral root (**Fig. 3A-C, Movie S4**). The arculum is then coupled to the base of the tibia-extensor tendon and to the tibia joint (**Fig. 3C**).

**Fig. 3.**
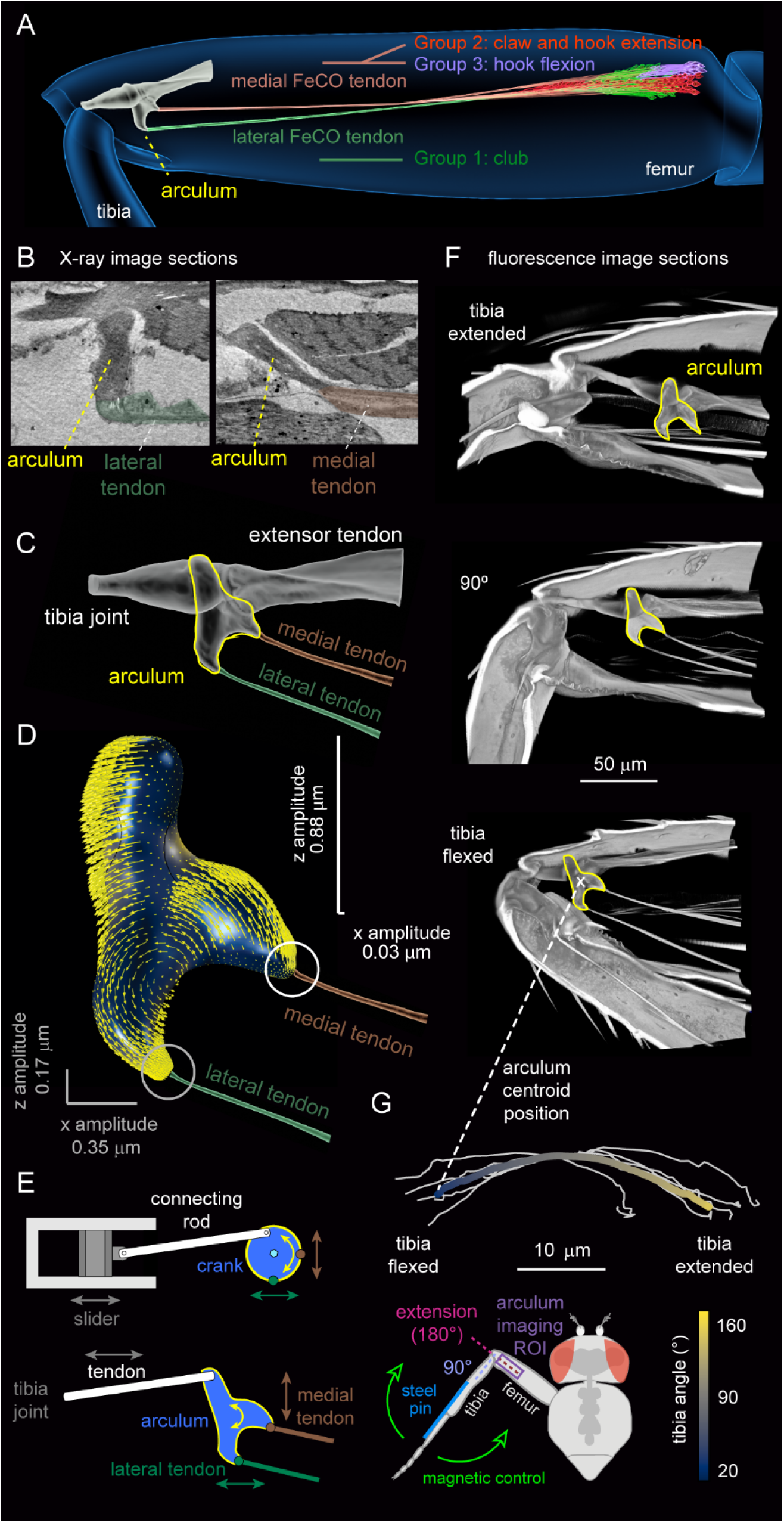
The arculum decomposes and transmits distinct mechanical signals to different subtypes of FeCO sensory neurons. **(A)** FeCO neurons are mechanically coupled to the tibia via two sensory tendons that converge upon the arculum. **(B)** X-ray image sections showing how the lateral and medial tendons attach to the arculum. **(C)** The arculum and its tendons, segmented from a confocal image of the femur. (**D**) A 3D finite element model of the arculum, stimulated by low amplitude, periodic forces at its base (top). Yellow arrows represent the arculum movement during vibration; the arculum rotates and the attachment point for the medial tendon (right) moves mainly in the z direction, while the attachment point for the lateral tendon (bottom) moves in the x direction. **(E)** A schematic showing how linear motion is translated into rotation in a manner analogous to a slider-crank linkage. (**F**) Autofluorescence images of the arculum at different femur-tibia joint angles. As the tibia flexes, the arculum translates toward the femur/tibia joint. A white “x” marks the position of the arculum centroid that we tracked with *in vivo* imaging in G. (**G**) Measurements of arculum position during tibia flexion/extension with trans-cuticular two-photon imaging (setup schematized below). The plot above shows arculum centroid position during full tibia flexion (white lines; n = 6 flies). The thick colored line is the average trace (color indicates the tibia angle).

Our finding that different sensory neuron subtypes are attached to the tibia via different tendons points toward a biomechanical origin for differences in mechanical feature selectivity. We did not observe efferent innervation of the FeCO, nor did we see connections between the FeCO and the surrounding muscles, as was reported in a previous study (Shanbhag et al., 1992). To investigate how tightly each type of sensory neurons are attached to the surrounding tissues, we expressed nuclear localized RFP in different types of FeCO neurons and used *in vivo* transcuticular two-photon imaging to track the location of the cell bodies while moving the tibia for its entire flexion/extension range with magnetically controlled setup (**Fig. S5A**). We found that the cell bodies of club and hook flexion cells move very little (< 2 μm) when the tibia is fully flexed and extended, suggesting that these cells are attached or embedded in stiffer tissue (**Fig. S5B, C; 4^th^ and 5^th^ row**). However, claw cells and hook extension cells moved distally towards the joint during tibia flexion, suggesting that these cells experience less resistance from the surrounding tissue (**Fig. S5B, C; 1^st^-3^rd^ row**). Interestingly, the translation of cells located on the distal side of the FeCO was greater than cells on the proximal side of the FeCO (**Fig. S5C**). These differences in cell translation further suggest that there may be differences in the mechanical forces experienced by different FeCO cells during tibia movement.

Overall, both our anatomical (**Fig. 1, 3A-C**) and physiological (**Fig. S2, Movie S3**) data point to the hypothesis that differences in FeCO sensory neuron activity are determined by differences in how mechanical forces are transmitted from the tibia via the arculum. We therefore sought to understand how the different tendons transmit mechanical forces to FeCO sensory neurons during flexion/extension movements of the femur-tibia joint, which is the major movement of this joint during walking in *Drosophila* (Karashchuk et al., 2021). We focused our analysis on the club and claw neurons because they attach to distinct tendons (lateral vs. medial), are selective for distinct kinematic features (vibration/movement vs. position), move very differently during tibia movements (stay immobile vs translate distally), and we possess clean genetic driver lines that specifically label these cell populations (**Fig. 1C-D, S2F**).

### The arculum decomposes and transmits distinct mechanical signals to different FeCO compartments

We first investigated how mechanical forces are transmitted during microscopic (< 1 μm), high frequency (100-1600 Hz) tibia vibrations; stimuli that excite club but not claw neurons (Mamiya et al., 2018). Because the lateral and medial tendons attach to different parts of the arculum (**Fig. 3C**), we focused on understanding how these two attachment points move during tibia vibration. Arculum movements during tibia vibration were too small to track *in vivo*, so we used a finite element model to investigate how the arculum responds to vibration (**Fig. 3D**). In the model, the arculum was suspended in the femur by four tendons (the tibia joint tendon that connects the arculum to the tibia, tibia extensor tendon, lateral FeCO tendon, and medial FeCO tendon). We modeled each tendon as a spring. The attachment points of the tendons were not rigid, allowing the arculum to both rotate and translate. To simulate joint vibration, we applied a periodic linear force to the tibia joint tendon. The model predicted that the arculum rocks in response to tibia vibration, causing it to rotate slightly as the tibia oscillates. This rocking moves the lateral and medial roots of the arculum in different directions (**Fig. 3D)**. Movements of the lateral root were larger along the long axis of the femur, while movements of the medial root were larger along the orthogonal axis (**Fig. 3D**). This result was consistent across a wide range of tibia vibration frequencies (**Fig. S6A**). The rocking of the lateral root was also visible at a larger scale in bright-field imaging of the arculum during spontaneous leg movements (**Movie S5).**

Our model suggests that the arculum decomposes the linear movement of the tibia into two orthogonal vectors. A helpful analogy for this decomposition is a slider-crank linkage, which is used in engines to convert linear motion into rotatory motion (**Fig. 3E**). In the case of the arculum, the two movement vectors are then transmitted along two different tendons: the lateral FeCO tendon transmits on-axis movements to the dendrites of the club cells, while the medial FeCO tendon transmits primarily off-axis movements to the claw neurons, reducing on-axis movements by as much as 30x. This difference between the two tendons, together with our observation that the club neurons are anchored more firmly to the surrounding tissues, provides a potential explanation for our prior observation that the mechanosensory threshold of claw neurons is more than 10 times higher than that of club neurons (Mamiya et al., 2018).

We next sought to understand how slower, nearly static, mechanical forces are transmitted to FeCO cells during macroscropic (1-50 µm) changes in femur/tibia joint angle, such as those that occur during self-generated tibia movements like walking (Karashchuk et al., 2021). When the tibia flexes, the arculum translates distally toward the femur/tibia joint (**Fig. 3F-G**). When the tibia extends, the arculum translates proximally, away from the femur/tibia joint. Thus, the primary effect of arculum translation is to pull/push the FeCO tendons along the long axis of the femur. This is consistent with our previous finding that large flexion/extension movements excite both claw and club neurons (Mamiya et al., 2018).

We took advantage of the autofluorescence of the arculum and FeCO tendons to visualize their movements during tibia flexion and extension with *in vivo* two- and three-photon imaging while passively moving the femur-tibia joint (**Fig. 3G**). The autofluorescence of the FeCO tendons was brighter under three-photon excitation (1300 nm); however, the faster repetition rate of the two-photon excitation laser (920 nm) was required to track arculum movements *in vivo* (**Fig. S6C, D**). Overall, our results from *in vivo* multiphoton imaging (**Fig. 3G)** were consistent with confocal images of the arculum in which the muscles and other soft tissues were digested for optical clarity (**Fig. 3F**). During tibia flexion, the arculum translates distally in a slightly curved trajectory (**Fig. 3G**). In addition to these translational movements, the arculum also rotates slightly to be more perpendicular to the long axis of the femur as the tibia flexes (**Fig. S6B**). As the arculum translates, the FeCO tendons do not bend, indicating that they are stiff. Therefore, we conclude that flexion and extension of the tibia causes the arculum and FeCO tendons to translate distally and proximally within the femur, which alters the mechanical strain on the FeCO dendrites.

In summary, our measurements of macroscopic (1-50 µm) arculum movement during tibia flexion/extension (**Fig. 3F-G**) and simulations of microscopic (< 1 µm) arculum movement during tibia vibration (**Fig. 3D-E**) suggest that this unique biomechanical structure is capable of transmitting distinct mechanical signals to distinct proprioceptor subtypes.

### Biomechanical mechanisms of position tuning among claw cells

As a population, claw neurons encode femur-tibia joint position, with specific cells increasing their activity at specific joint angles (Mamiya et al., 2018). However, it remains unclear how different claw cells achieve their angular feature selectivity. We found above that the arculum moves back and forth within the femur as the tibia flexes and extends (**Fig. 3F-G**), which we propose increases and decreases tension on the claw cell dendrites via the medial tendon. One possible mechanism for position tuning among claw neurons is that the mechanical coupling between the medial tendon and the claw cells distributes a different level of strain to each proprioceptor neuron. Because directly measuring strain within the FeCO is technically infeasible, we took a four-step experimental and modeling approach to test if this mechanism is consistent with the structure and dynamics of the proprioceptive organ (subheadings i-iv, below). We first experimentally measured the movement of claw neurons and associated cap cells during tibia movement and following transection of the tendons. Second, we used the X-ray reconstruction to make a mechanical model of the claw cells and the medial tendon, to test if the anatomy of the system is sufficient to explain measured patterns of claw cell movement. Third, we used this minimal model to predict how strain is distributed among the claw neurons when the tibia is moved. Fourth, we tested model predictions using *in vivo* calcium imaging from claw proprioceptors in the fly leg during tibia flexion/extension.

### (i) Claw neurons and cap cells are held under resting tension and translate within the femur during tibia movement

We fluorescently labeled claw neurons and their cap cells and tracked their movement within the femur using *in vivo* two-photon imaging during passive movements of the femur-tibia joint (**Fig. 4A**). Like the arculum, claw neurons and their cap cells translate distally when the tibia flexes and proximally when the tibia extends (**Fig. 4B, C**). In addition to the large movements along the distal-proximal axis of the femur, claw neurons and cap cells also translate slightly along the perpendicular axis (**Fig. 4B, S7C**). Interestingly, we also noticed that the distance that claw cell bodies and cap cells translate during tibia flexion and extension increases linearly with their relative position along the proximal to distal axis of the femur (**Fig. 4E, S8A, C**). In other words, distal claw neurons and cap cells move farther than proximal cells for a given change in tibia angle. These systematic differences suggested to us that the strain on claw neurons during tibia flexion/extension might depend on their position within the array.

We next asked whether the medial tendon applies tension to claw neurons throughout the entire range of tibia angles. Alternatively, there could exist an equilibrium angle where the tension becomes zero, beyond which the tendon compresses the neurons. To distinguish between these two possibilities, we cut the tendons that connect the arculum to the FeCO while holding the tibia at different angles (**Fig. S9A**). We then tracked the resulting displacement of claw cap cells (**Fig. S9A, B)**. We found that the cap cells relaxed proximally within the femur after the tendons were cut, regardless of the tibia angle. The distance the cap cells relaxed was greater when the tibia was flexed prior to cutting (**Fig. S9B**). Consistent with the movements of the cells we observed during the tibia flexion/extension, the relaxation distance of cap cells increased linearly with their relative position along the proximal to distal axis of the femur (**Fig. S9B, C)**. In summary, these results indicate that the claw cells are held under tension, even when the tibia is fully extended, and this tension increases as the tibia is flexed.

### (ii) A biomechanical model of claw neurons shows that cell movement patterns can be explained by the geometry and material properties of the organ

To test whether the geometry and the material properties of the elements in the FeCO are sufficient to explain the observed cell movements (**Fig. 4E**), we constructed a series of finite element models. We based the model on the geometry of the FeCO and tendon attachments, measured from the X-ray reconstruction (**Fig. 1**), and the tension we observed in the FeCO (**Fig. S9**). The model consisted of a series of coupled mechanical elements: claw cell bodies, dendrites, cap cells, tendon fibrils, the medial tendon, and the arculum (**Fig. 4D**). We modeled these elements as linear elastic materials and embedded them in elastic tissue mimicking the extracellular matrix surrounding the FeCO (see Methods for details). When we moved the tendons according to measured trajectories of the arculum during tibia flexion/extension (**Fig. 3G**), the model effectively reproduced the trajectories of claw and cap cell movements during tibia extension and flexion (**Fig. 4F, S7B, S8B, D**). For example, distal cells translated farther than proximal cells during tibia flexion (**Fig. S8B, D**), as we saw in measurements of claw and cap cell movement (**Fig. 4E, S8A, C**). The model results suggest that these dynamics arise from the basic geometry of the FeCO and tendon attachments, and the material properties of these elements.

Because it is not possible to measure mechanical strain on claw neurons *in vivo,* we used the model to predict how strain on these cells changes during tibia flexion/extension. The model predicted a gradient of strain among claw neurons in which the strain was always higher for more proximal cells than more distal cells (**Fig. 4G, H, Movie S6**). The difference in the strain became larger as the tibia was flexed (**Fig. 4G, H, Movie S6**). These results are consistent with the hypothesis that the medial tendon distributes a different level of strain to each claw cell.

Which properties of the FeCO are important for establishing the gradient of cell movement and mechanical strain among claw cell dendrites? By modifying different components of the model (**Fig. S10**), we were able to identify three key factors. The first factor was the orientation of the fibrils that connect the array of claw cap cells to the medial tendon, which we measured from the X-ray reconstruction (**Fig. S10A)**. Because the fibrils fan out from a single endpoint of the medial tendon (**Fig. S10C**), the fibrils that connect to more proximal claw cells are oriented more parallel to the direction of tendon movement, while those that connect to more distal claw cells are more perpendicular (**Fig. 4D**). When the tendon is pulled toward the joint during the tibia flexion, the fibrils connected to more proximal cells transfer most of the force in a direction that stretches the proprioceptor dendrites and increases the strain in these cells (**Fig. 4G, H**). Results from modeling experiments in which we manipulated the fibril angles by moving the fibril branching point suggest that the fibrils oriented closer to the movement of the tendon transfer more strain to the dendrites (**Fig. S10D**). The second factor was the stiffness of the tissues surrounding the cells, which we were unable to measure directly, but for which published values exist (Levental et al., 2007). Modelling experiments showed that stiffer tissues reduced the cell movements, while softer tissues enhanced cell movements (**Fig. S10E**). The third factor was the stiffness of the fibrils and dendrites relative to the surrounding tissues, which we were also unable to measure directly. Stiffer fibrils and dendrites transfer forces more effectively through the softer surrounding tissues, and cells moved more when we made these elements stiffer (Fig. S10F). The effect of the fibrils/dendrite stiffness on the dendritic strain was more complex because the dendritic strain depends on the relative motion between the claw cell and the connected cap cells. We found that the strain gradient is maintained as long as the fibrils and dendrites had a stiffness close to published values (Frantsevich et al., 2019; Gosline et al., 2002; Rydholm et al., 2010) or stiffer, but the gradient reversed when the fibrils and dendrites were much softer (Fig. S10F). Overall, these results suggest that the basic geometry of the fibrils and the stiffness of the elastic elements are sufficient to generate the differential movements of claw cells and a strain gradient in their dendrites.

### (iii) The finite element model predicts the existence of a goniotopic map of joint angle among claw proprioceptors

The gradient of dendritic strain made a further intriguing prediction: the existence of a map of leg joint angle across the linear array of claw cells. For example, if all flexion selective claw neurons had similar sensitivity to the increase in the mechanical strain, proximal neurons will become active first as the tibia flexes, followed by more distal neurons. We define a new term to describe this map: *goniotopic* (*gōnia* is the Greek word for angle).

The model made two additional predictions about the orientation and robustness of the goniotopic map. Because strain on the claw cell dendrites increases as the tibia flexes and decreases as the tibia extends (**Fig. 4G, H**), the model suggests that the flexion selective claw neurons detect increases in strain while the extension selective claw neurons detect decreases in strain. This means that the orientation of the extension map is the reverse of the flexion map. As the tibia extends past 90 degrees, distal claw neurons should become active first, followed by more proximal neurons. Due to smaller difference in strain among claw neuron dendrites during tibia extension, the model also predicts that the extension map should be noisier and less robust than the flexion map.

### (iv) Claw proprioceptors in the fly leg contain a goniotopic map of leg joint angle

To test for the existence of a goniotopic map of leg joint angle among claw neurons, we developed new methods to measure the angular tuning of claw cell bodies inside the femur with volumetric two-photon calcium imaging during passive movements of the tibia. Imaging somatic GCaMP signals together with activity-independent tdTomato fluorescence allowed us to track individual cells and simultaneously measure neural activity, relative cell position, and claw cell movement (**Fig. 5A, Movie S7**). Consistent with the finite element model prediction, we found that flexion of the tibia first activated flexion-tuned claw neurons in the proximal FeCO, followed by the activation of increasingly distal cells at more acute angles (**Fig. 5A-C, Movie S7**). This goniotopic map was consistent across flies and different genetic driver lines labeling claw flexion neurons (**Fig. 5C**, **S11 1^st^ and 2^nd^ columns**).

**Fig. 5.**
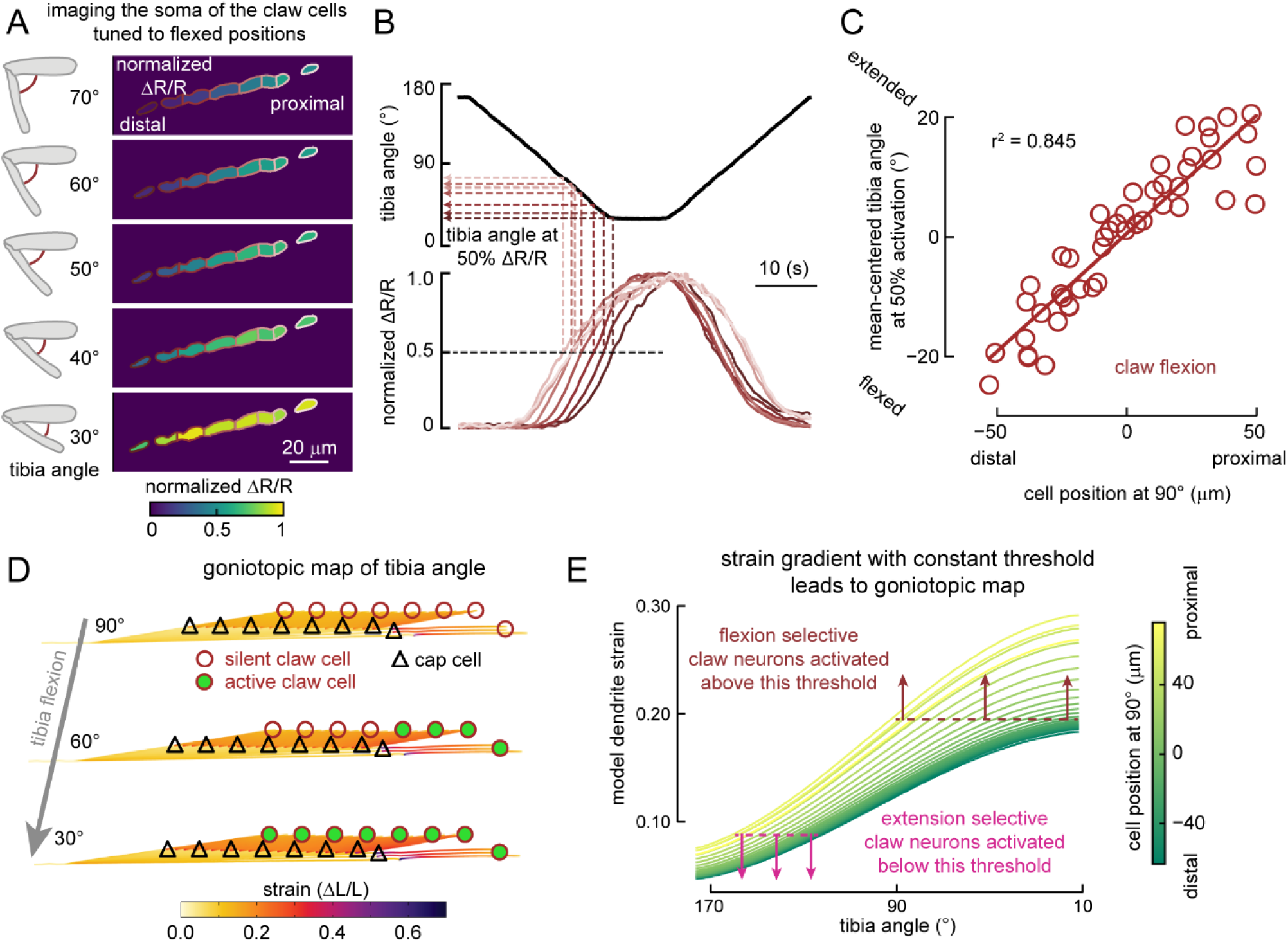
A goniotopic map of tibia joint angle among claw neurons. **(A)** Example images of GCaMP7f fluorescence (left column) and normalized activity (right column) of flexion-selective claw neurons during slow tibia flexion (6°/s) recorded with two photon microscopy. Each colored circle indicates a tracked cell. Claw cell bodies move distally and increase their calcium activity as the tibia flexes. (**B**) Tibia angle (top) and normalized calcium activity (bottom) during the example recording shown in **A**, indicating the tibia angle where each cell reached 50% of its peak activity. The color scheme is the same as in **A**. More distal cells (darker shades of copper) reached 50% of their peak activity at more acute angles. (**C**) A linear relationship between each claw cell’s position along the proximal-distal axis of the femur and the tibia angle when the cell reached 50% of its maximum activity. Within each fly, we subtracted the mean of the tibia angle at which cells reached their 50% maximum activity (see Methods for details). (**D**) Schematic illustrating how the goniotopic map of tibia angle is represented by the activity of the flexion selective claw cells. **(E)** Illustration of how the strain gradient predicted by the finite element model combined with a uniform threshold for the flexion and extension selective claw neurons lead to a goniotopic map of tibia angle represented by the activities in the array of claw neurons. Strain plot is the same as the one shown in **Fig. 4H**. Dotted lines represent hypothetical thresholds for the activation of the flexion and extension selective claw neurons.

We observed a similar, though less robust, goniotopic map within the extension-tuned claw neurons (**Fig. S11 3^rd^ and 4^th^ columns)**. Consistent with the prediction of the model, the extension map had the reverse orientation of the flexion map. Extension of the tibia first activated more distal claw extension neurons, followed by activation of more proximal neurons as the tibia extended further (**Fig. S11 3^rd^ and 4^th^ columns**). The difference in the robustness of the two goniotopic maps may be due to the smaller differences in strain experienced by different claw neurons during extension, as predicted by the model (**Fig. 4H**).

In summary, our measurements of claw neuron movement and calcium activity reveal a topographic map of joint angle across the linear array of claw neurons (**Fig. 5D, E**). Flexion and extension selective claw neurons appear to be intermingled in this array (**Fig. S11**). For flexion selective claw neurons, proximal neurons are active first at more obtuse angles during tibia flexion (< 90°), and distal neurons are active later at more acute angles. For extension selective claw neurons, distal neurons are active first at more acute angles during tibia extension (> 90°), and proximal neurons become active at more obtuse angles. Combined with the model predictions of an equivalent map of strain accumulation, these results point to a biomechanical origin for joint angle feature selectivity among claw neurons (**Fig. 4G, H, 5E**).

### Club neuron dendrites contain a tonotopic map of tibia vibration frequency

Finally, we investigated whether a topographic map exists among club neurons. Unlike claw neurons, which translate within the femur as the tibia flexes and extends, club neurons move very little (< 2 μm; **Fig. S5C; bottom row**), suggesting that the tissues surrounding club cells may be stiffer than those surrounding claw neurons (**Fig. S10E**) or that club neurons are anchored to the femur cuticle. Immobilizing cell bodies may contribute to club neuron sensitivity by increasing mechanical strain between club cells and the tendon during tibia movement.

To test for the presence of a tonotopic map within the FeCO, we vibrated the tibia at different frequencies (100-1600 Hz; 0.9 μm amplitude) and recorded calcium activity from club neurons in the femur with two-photon imaging (**Fig. 6A**). During vibration, we saw consistent increases in intracellular calcium in the dendrites and cell bodies of the club neurons. Because the activity in the dendrites were larger, presumably due to calcium currents generated by mechanosensory channels, we focused on the dendritic activity (**Fig. 6B-C**). As we increased the vibration frequency, the response amplitude increased and reached a plateau around 800 Hz (**Fig. 6C**). In addition to the increase in response amplitude, we also observed a shift in the spatial distribution of calcium activity (**Fig. 6B, D**). As the vibration frequency increased (200-1600 Hz), the center of the response moved from the distal/lateral side of the FeCO to the proximal/medial side (**Fig. 6B, D**). This shift was consistent and statistically significant across flies (**Fig. 6D, S12**). Thus, the FeCO also contains a tonopic map of tibia vibration frequency within the dendrites of the club neurons, although the mechanism that establishes this map remains unclear (see Discussion).

**Fig. 6.**
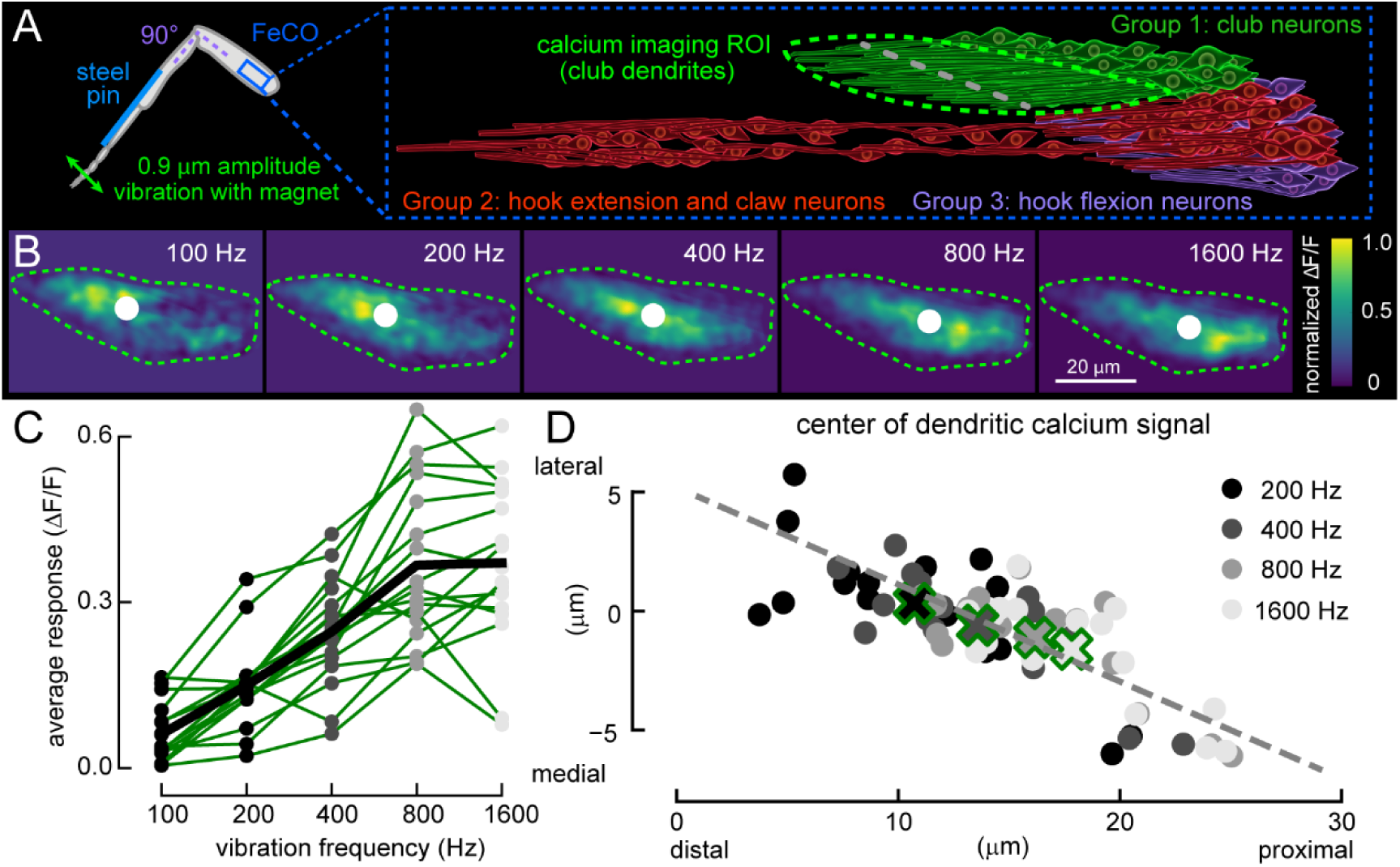
A tonotopic map of tibia vibration frequency among club neurons. **(A)** Schematic of calcium imaging from the club neuron dendrites during tibia vibration using transcuticular two-photon microscopy (left). **(B)** Examples of calcium activity (ΔF/F, normalized for visualization) in the club dendrites during vibration of the tibia. White circles represent the weighted center of the calcium activity. Green dotted lines show ROI of club dendrites. **(C)** Average calcium activity (ΔF/F) in the club dendrites of individual flies (n= 17 flies) during the vibration of the tibia at different frequencies. Circles and thin lines are from individual flies and the thick black line represents the average across flies. **(D)** A scatter plot showing the weighted center of calcium activity for each vibration frequency (200 to 1600 Hz; n = 17 flies). Larger X’s with green outlines indicate averages. The gray line shows the best-fit for the response centers and is also indicated in **A** to illustrate the primary spatial axis of the tonotopic map.

## Discussion

We investigated two non-mutually exclusive mechanisms for establishing mechanical feature selectivity among proprioceptors in the *Drosophila* leg. One possibility is that each proprioceptor subtype uses different mechanosensensory or other ion channels, allowing sensory neurons to respond selectively to different kinematic features (e.g., joint position, directional movement, vibration). Another possibility is that biomechanical elements transmit different mechanical forces to each proprioceptor subtype. Our RNA-seq data suggest that there are not large differences in the expression of mechanosensory or voltage-gated ion channels among different subtypes of FeCO neurons (**Fig. 2, S3, S4**). On the other hand, we identified two biomechanical mechanisms that could allow specific subtypes of FeCO proprioceptors to detect distinct joint kinematic features (summarized in **Fig. S13**).

First, a biomechanical model based on anatomical reconstruction of the fly leg predicts that the arculum transmits distinct aspects of tibia joint movement to the FeCO via distinct tendons (**Fig. 3, S13**). The two tendons move similarly for macroscopic (1-50 µm), low frequency changes in tibia angle, such as those that occur when a fly walks. However, during microscopic (< 1 µm), high frequency vibrations of the tibia, the shape of the arculum allows it to push and pull the lateral tendon in a direction that excites the club sensory neurons, while moving the medial tendon in a perpendicular direction that does not excite the claw sensory neurons. This model of the arculum’s function is consistent with all of our anatomical and physiological measurements. However, a direct test of this model must await the development of new techniques to measure sub-micron movements of the arculum during natural joint movements.

Second, a finite element model of the claw neurons predicts the existence of a strain gradient across position-tuned proprioceptors. The geometry of the model is taken directly from anatomical reconstruction of the FeCO and attachment structures from X-ray tomography (**Fig. 1**). It reconciles three key experimental results. First, tracking the movement of claw neurons during tibia joint movement revealed that distal claw neuron cell bodies translate more during tibia flexion/extension compared to proximal cell bodies (**Fig. 4**). Second, recordings of calcium signals from proprioceptor cell bodies during joint movement revealed a goniotopic map of leg joint angle among flexion and extension tuned claw neurons (**Fig. 5**). Third, severing the medial tendon causes the claw neurons and cap cells to retract proximally (**Fig. S9**), which indicates that claw neurons are held under resting tension. Without this resting tension, the goniotopic map is disrupted. Further analysis of the finite element model revealed that the gradient in the angle of the medial tendon fibrils, the stiffness of the tissues surrounding the cells, and the stiffness of the fibrils and dendrites relative to the surrounding tissues, are all important for establishing the movements of the claw neurons and the gradient of strain in the claw neuron dendrites **(Fig. S10).** Overall, this model is consistent with all of our anatomical and physiological measurements of position-tuning in claw neurons. Again, new techniques to measure mechanical forces on the proproprioceptor dendrites in the intact leg would be needed to directly test this model.

A remaining mystery is how claw neurons sense tibia extension. The gradient of dendritic strain in the finite element model suggests that the extension selective claw neurons are responding to a decrease in dendritic strain (**4G, H, 5E**). Sensing decreases in strain is difficult to reconcile with the force-from-lipid principle of mechanosensation, in which channels are opened through an increase in the force on the lipid bilayer (Teng et al., 2015). However, it may be consistent with the force-from-filament principle, in which mechanosensory channels are tethered to the cytoskeleton or extracellular matrix (Katta et al., 2015). An example of the force-from-filament principle is the tip link of mammalian hair cells, which contributes to the directional selectivity of the mechanosensory channels (Gillespie and Müller, 2009). If a similar tether exists between the mechanosensory channels in the dendrites of the claw neurons and the cap cells, it should be possible to control whether the channels open when tensile strain on the dendrites decreases or increases by switching the configuration of the tether. Thus, extension and flexion selective claw neurons may have different tethering configurations of mechanosensory channels, in order to respond to decreases or increases in dendritic strain, respectively.

We also identified a map of tibia vibration frequency among club neurons (**Fig. 6**). However, anatomical reconstruction and RNA-seq of FeCO nuclei did not resolve the mechanism of this tonotopic map. One possibility we considered is that the difference in the length of the tendon connected to each club cell could produce a different resonant frequency for each cell. However, a simple mechanical model based on measurements of the tendon fibrils from the X-ray microscopy data did not support this hypothesis: neither the longitudinal nor transverse vibrational modes would be sufficiently different to produce large enough changes in resonance at the club neuron dendrites. Another possibility is that systematic differences in the stiffness of each club neuron dendrite or in the mass of the cap cell could underlie their frequency tuning, similar to the situation in katydid hearing organs (Hummel et al., 2017). Finally, the tonotopic map could be established through tuning based on electrical resonance within club neurons, as occurs within the vertebrate cochlea (Fettiplace and Fuchs, 1999). Again, the development of techniques to directly measure dendritic strain from FeCO neurons would help to distinguish between these alternative mechanisms.

Our RNA-seq results do not support the hypothesis that different types of FeCO neurons express distinct types of mechanosensory channels or voltage-gated ion channels (**Fig. 2, S3, S4**). However, because our analyses cannot detect splice variants or post-translational modifications of the channels, it is still possible that some of the feature selectivity of FeCO neurons are determined by the type of mechanosensory or voltage gated sodium/potassium channels these cells express. This could be especially true for the differences in the feature selectivity within a group of neurons that appear to have minimal differences in biomechanical filtering, such as the frequency turning differences among the club, or differences in the responses between the flexion and the extension tuned claw cells. Alternative splicing of the *slo* gene, which encodes a voltage-gated potassium channel, is thought to underlie the gradient of frequency tuning in the vertebrate cochlea (Fettiplace and Fuchs, 1999). We found that *slo* is highly expressed in FeCO neurons (**Fig. S4**), which raises the possibility that a similar mechanism underlies frequency tuning in club neurons.

One proprioceptor subtype that we did not address in detail in this paper is the hook neurons. Hook neurons are either flexion or extension sensitive (**Fig. 1D, S2F**). These two types seem to receive distinct mechanical forces, based on their attachments to the tendons and surrounding tissues. Extension selective hook neurons are located near the distal edge of the FeCO, attached to a branch of medial tendon that connects to the claw neurons, and translate distally during tibia flexion (**Fig. 1, S1B, S2F, S5**). Flexion selective hook neurons are located closer to the proximal edge of the FeCO, attach to a separate branch of the medial tendon, and do not translate when tibia was flexed (**Fig. 1, S1B, S2F, S5**). As with the tonotopic map in the club neurons, direct measurements of dendritic strain may be necessary to understand mechanisms of feature selectivity in hook neurons.

Our biomechanical model did not include scolopale cells, a cell that surrounds the dendrites of chordotonal cells, because their biomechanical properties and function are not clear (**Fig. 1**). In all chordotonal organs, across species, each scolopale cell surrounds the dendrites of chordotonal neurons that are attached to the same cap cells (Field and Matheson, 1998). One consequence of the wrapping of the dendrites by scolopale cells may be an interference between the activities of the neurons that share the same extracellular space, similar to the inhibition among *Drosophila* olfactory sensory neurons sharing the same sensillum (Su et al., 2012). Better methods for identifying the pair of neurons that connect to the same cap cell *in vivo* will be necessary before we can investigate the possible interactions between the pair of FeCO neurons wrapped by a scolopale cell.

In the current study, we passively moved the tibia to better control its position and velocity. Although we did not see any connections between the FeCO neurons and the surrounding muscles, the tibia extensor muscle is connected to the arculum (**Fig. 3C**). This raises the possibility that the mechanical forces experienced by FeCO neurons during active movements of the tibia may be different from those experienced during passive movements. Although measurements of the forces in the FeCO during active and passive movements would be needed to resolve mechanical differences, these differences may not be large, because the responses of the club and claw neurons are similar between active and passive movements of the leg (Dallmann et al., in prep).

Topographic neural maps are found throughout the animal kingdom, including in other peripheral mechanosensory organs. For example, hair cells in the vertebrate cochlea (Fettiplace and Fuchs, 1999) and chordotonal neurons in the katydid tympanal organ (Oldfield, 1982) are organized into tonotopic maps of auditory frequency, similar to club neurons in the fly FeCO. Fractionation of the tibia joint angle range across position-tuned proprioceptors has been previously described in the locust FeCO (Field, 1991; Matheson, 1992). Translation of FeCO cell bodies during limb movements has also been observed in other insects (Nishino et al., 2016; Nishino and Sakai, 1997). However, to our knowledge, goniotopic maps have not been described previously, perhaps due to the difficulty of recording activity from populations of mobile proprioceptive sensory neurons embedded within the body. Our findings that simple geometry and material properties are enough to generate goniotopic map raise the possibility that goniotopic maps of proprioceptive stimuli may exist in other organisms and sensory organs, thus creating a scaffold for a central topographic representation of the body.

Are the maps established in the fly leg preserved within central circuits? Similar to the tonotopic map we describe here among club neurons, there exists a map of vibration frequency among club axons in the VNC (Mamiya et al., 2018). Downstream VNC interneurons are also tuned to specific frequency ranges, indicating that they sample selectively from the topographic projections of club axons (Agrawal et al., 2020). In comparison, calcium imaging from position-tuned axons failed to resolve any topographic organization (Mamiya et al., 2018). However, interneurons downstream of claw axons encode specific joint angle ranges (Agrawal et al., 2020; Chen et al., 2021), suggesting that the goniotopic map may be preserved at the level of connectivity. Elucidating this organization will require reconstruction at the synaptic level, for example using connectomics of VNC circuits (Azevedo et al., 2022; Phelps et al., 2021).

## Acknowledgements

We thank members of the Tuthill laboratory for technical assistance and feedback on the manuscript, particularly Ellen Lesser, who initially observed that FeCO neurons move within the leg, and Tony Azevedo, who was involved with the initial identification of the arculum within the x-ray dataset. We thank Thedita Pedersen for reconstructing FeCO neurons and dendrites from the X-ray microscopy dataset. We thank Katie Stanchak and Bing Brunton for discussions and helpful comments on the manuscript. We thank Bertil Hille for suggesting the term *goniotopic*. We thank Rachel Wong and Gwyneth Card for generous sharing of equipment. We thank Wyeth Bair and the members of the Bair laboratory for sharing their 3-photon microscope and related equipment, which were purchased through support from NIH grants (R01 EY010115 and R34 NS116733) to Wyeth Bair, and the support from NIH (supplement to R01 EY 018839) and National Primate Research Center (supplement to P51OD01042559) to Anitha Pasupathy. We thank Jack Waters and Kevin Takasaki for assistance with the construction of the 3-photon microscope. We used stocks obtained from the Bloomington *Drosophila* Stock Center (NIH P40OD018537). We acknowledge support from the NIH (S10 OD016240) to the Keck Imaging Center at UW, and the assistance of its manager, Nathaniel Peters. We acknowledge ESRF for granting beamtime for the experiments: LS2705, LS2845, IHLS2928, IHLS3121, IHHC3498, IHMA7 and IHLS3004. This work was supported by the Visiting Scientist Program at HHMI/Janelia to C.C. and J.C.T, as well as a UW Innovation Award, a Searle Scholar Award, a Klingenstein-Simons Fellowship, a Pew Biomedical Scholar Award, a McKnight Scholar Award, a Sloan Research Fellowship, the New York Stem Cell Foundation, and NIH grants R01NS102333 and U01NS115585 to JCT, and NSERC Discovery (Grant no. 687216), early career supplement (675248), and an NSERC Canada research chair (Grant no. 693206) to NM, and funding from NIH (R01NS108410), and awards from the Edward R. and Anne G. Lefler Center and the Goldenson Family to W.C.A.L.. H.L. is a CPRIT Scholar in Cancer Research (RR200063), and supported by NIH (R00AG062746), the Longevity Impetus Grant, and Ted Nash Long Life Foundation. A.P. has received funding from the European Research Council under the European Union’s Horizon 2020 Research and Innovation Programme (grant no. 852455). JCT is a New York Stem Cell Foundation – Robertson Investigator.

## Author contributions

Conceptualization, A.M., P.G., N.M., and J.C.T; Methodology, A.M., P.G., I.S., J.S.P., A.S., Y. Q., T. L., A.P., A.K., N.M.; Investigation, A.M., P.G., I.S., Y. Q., T. L., J.S.P., A.S., N.M.; Resources, C.C., J.S.P, A.P., A.K, W.A.L. and H. L.; Analysis, A.M., P.G, I.S., Y. Q., T. L., A.S., N.M., J.C.T..; Writing, A.M., N.M., J.C.T.; Supervision, W.A.L., H. L., N.M., and J.C.T.; Funding Acquisition: W.A.L., H. L., N.M., and J.C.T.

## Supplemental Figures

**Fig S1.**
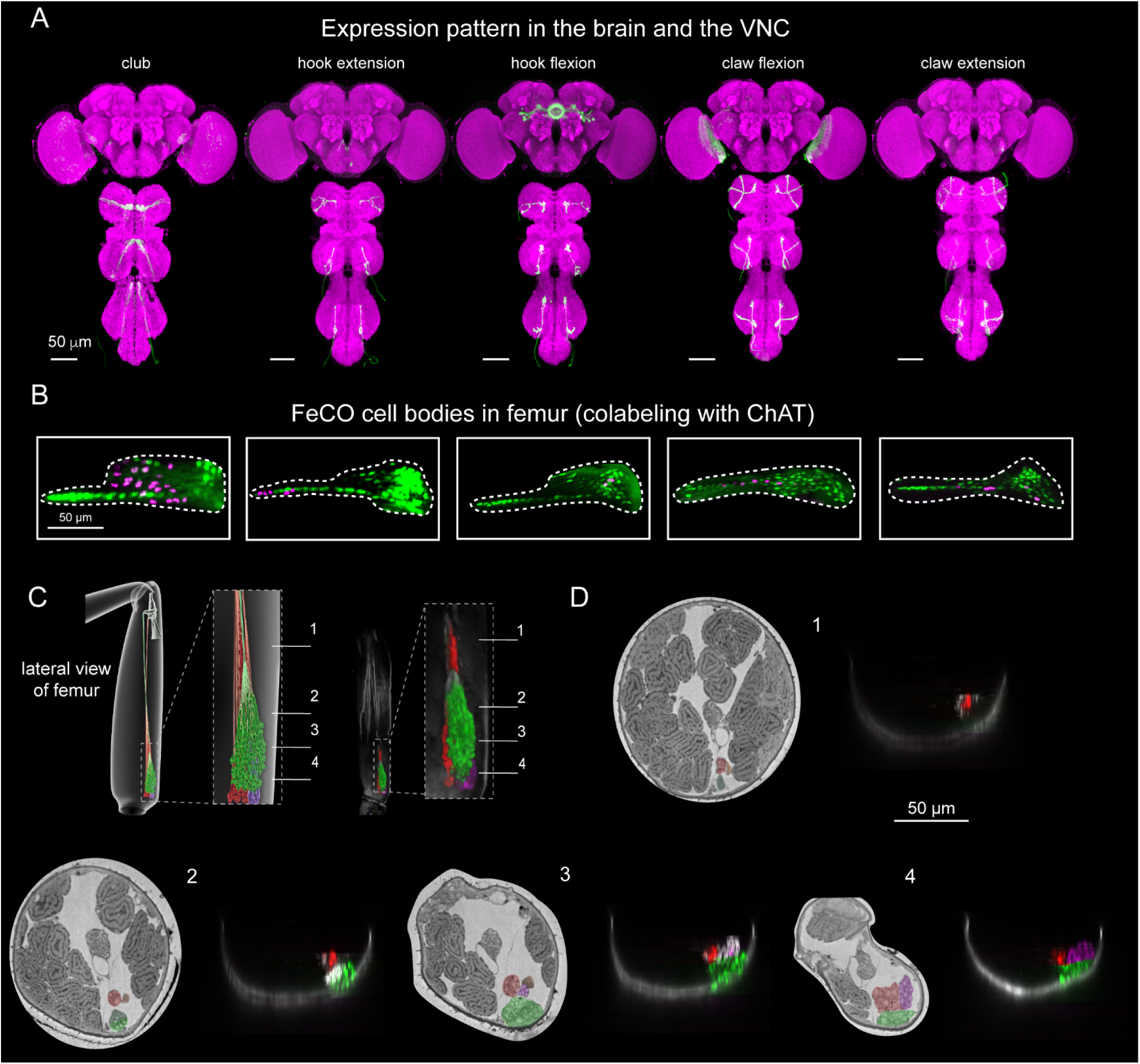
Proprioceptor subtypes are organized into functional compartments within the fly femur. **(A)** Confocal images of the fly brain and the VNC, showing the expression patterns of split-Gal4 drivers used to label specific subtypes of FeCO neurons (green; genotype listed in supplemental table 1). Magenta is a neuropil stain (nc82). **(B)** Confocal images of the FeCO cell bodies in femur, showing the expression pattern of Gal4 drivers used to label specific subsets of FeCO neurons (magenta). Green represents the expression pattern of ChAT-Gal4, which labels all mechanosensory neurons. **(C)** Same as Fig **1D**, but for cross sections of the femur. (**D**) Reconstruction of FeCO compartments from X-ray and confocal imaging of the fly femur. Each image corresponds to a transverse section indicated in the schematic in **C**. FeCO compartments and tendons are indicated by color shading.

**Fig S2.**
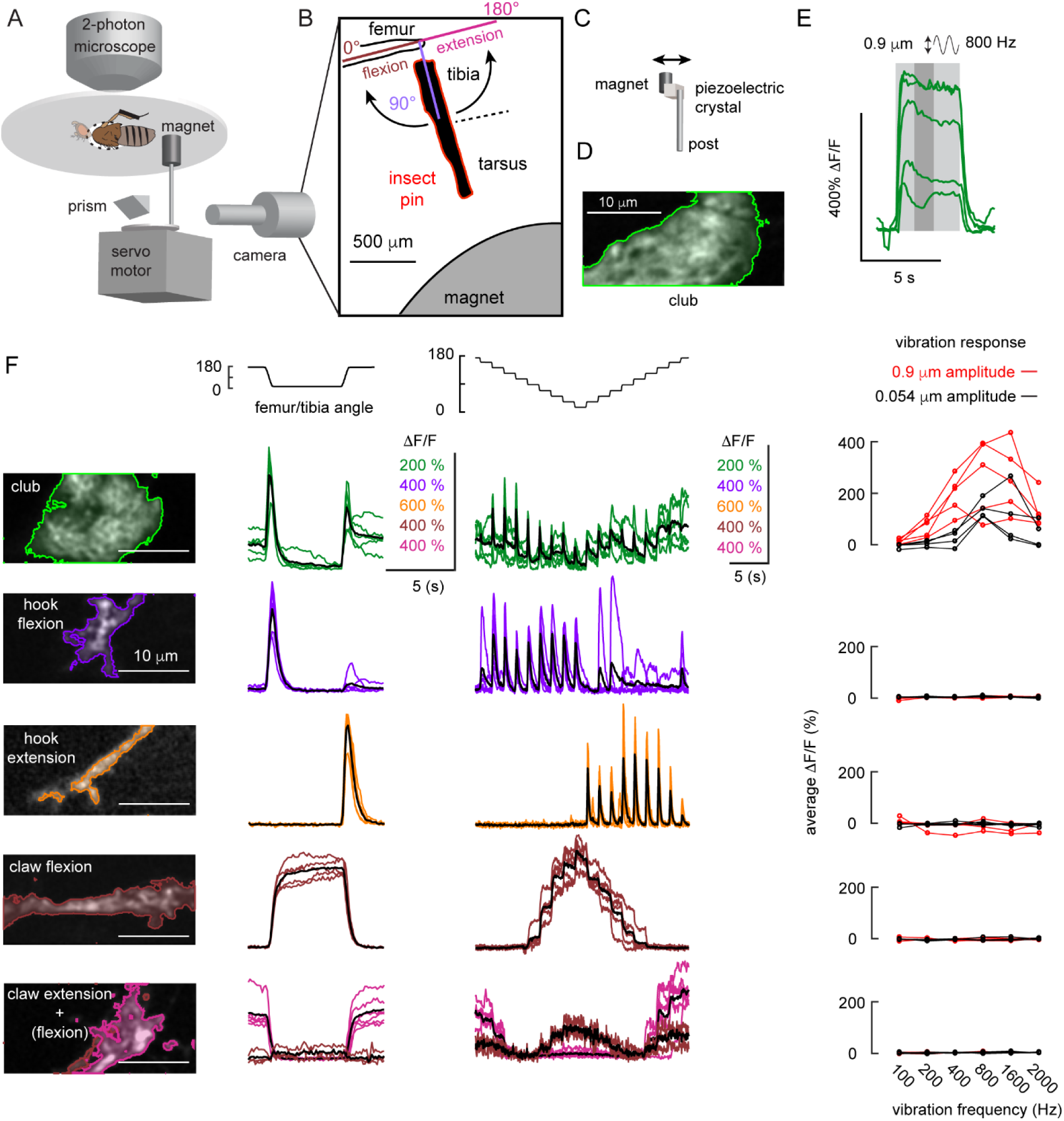
Calcium imaging from axons of FeCO split-Gal4 lines. **(A)** An experimental setup for two-photon calcium imaging from the axons of different types of FeCO neurons in the VNC while changing the femur-tibia joint angle. We controlled the joint angle by gluing a pin to the tibia and positioned it using a magnet mounted on a servo motor. We backlit the tibia with an IR LED and used a prism to record the view from the bottom with high-speed video camera. **(B)** An example frame from a video used to track joint angle. We painted the pin black to enhance the contrast against the bright background. **(C)** An experimental setup for vibrating the tibia with a magnet attached to a piezoelectric crystal. **(D)** An example two-photon image of the club axon terminal where we recorded the responses to the vibration stimuli (region of interest shaded with light green and surrounded by a green line). **(E)** Time courses of the club neuron’s response (ΔF/F) to an 800 Hz, 0.9 μm vibration of the magnet (n = 5 flies). Each line represents the response from one fly. We averaged the activity in a 1.25 s window (indicated by a dark gray shading) starting from 1.25 s after the stimulation onset and used it as a measure of response amplitude in the rightmost column of F. **(F)** Calcium responses (ΔF/F) of the axons of different subsets of FeCO neurons to swing (2^nd^ column), ramp-and-hold (3^rd^ column), and vibration (4^th^ column) stimuli. 1^st^ column shows the GCaMP fluorescence of the axons of different subsets of FeCO neurons. 1^st^ row: club (n = 5 flies), 2^nd^ row: hook flexion (n= 5 flies), 3^rd^ row: hook extension (n = 5 flies), 4^th^ row: claw flexion (n= 5 flies), 5^th^ row: claw extension and flexion (n= 6 flies). Because the Gal4 line for the 5^th^ row contained a small number of flexion selective claw neurons in addition to the extension selective claw neurons, we separated the pixels into two clusters based on their response similarity to the swing or ramp-and-hold stimuli using k-means clustering. Each trace is the average response from one fly.

**Fig. S3.**
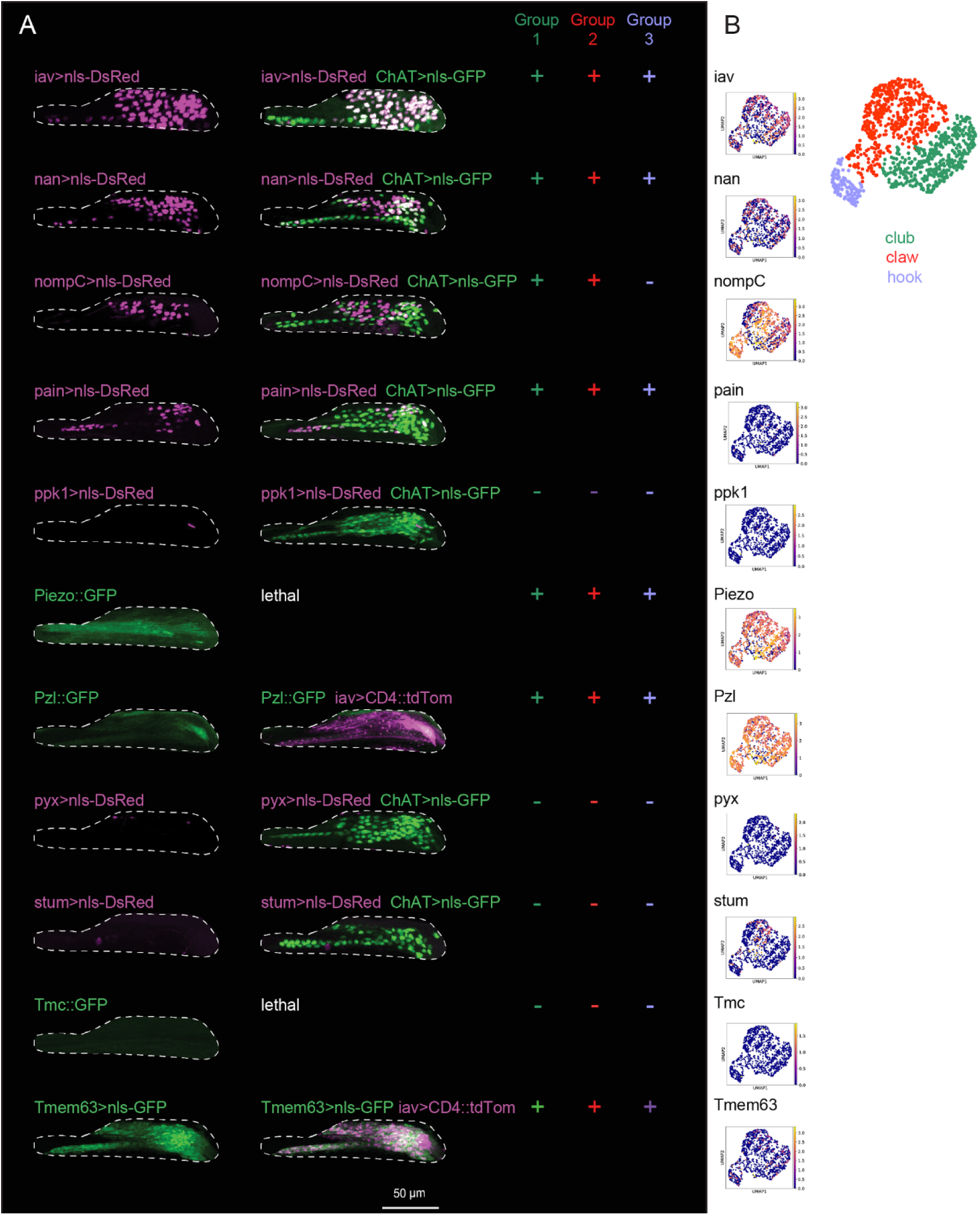
Mechanotransduction channel expression in the femoral chordotonal organ (FeCO). **(A)** Confocal images of mechanotransduction channel reporter expression in the FeCO cell bodies in the femur. Images are co-labeled with either ChAT- Gal4 (labels all mechanosensory neurons) or iav-Gal4 (labels most FeCO cell bodies). In some cases, co-labeled genotypes are lethal and not shown. (+) and (-) indicate expression within scoloparia groups 1, 2, and 3. **(B)** RNA transcript expression levels of the same mechanotransduction channels shown in A, displayed for the tentative club, claw, and hook neuron clusters identified in the RNA-seq dataset.

**Fig. S4.**
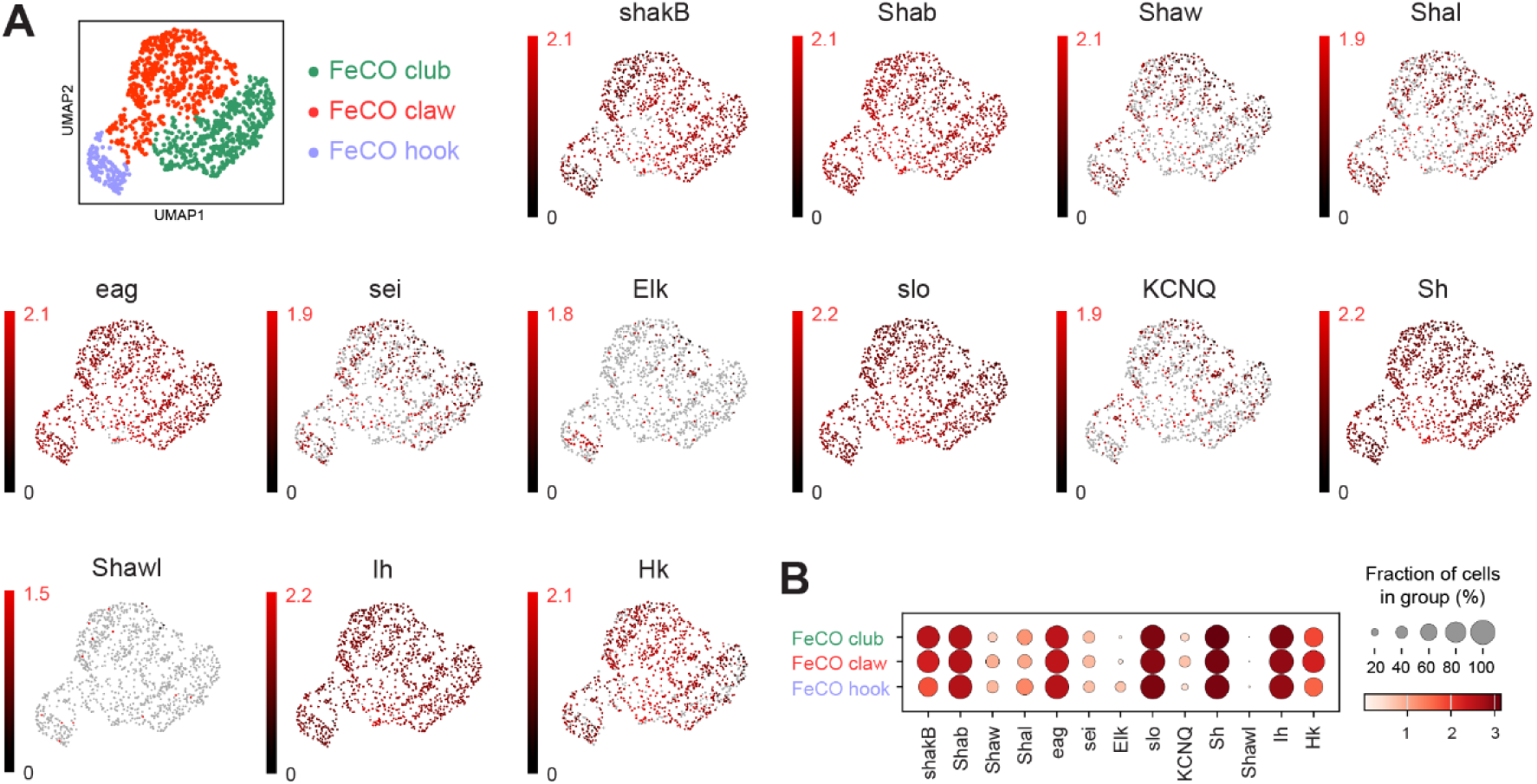
Single-nucleus RNA-sequencing reveals overlapping expression of voltage-gated channels among FeCO subtypes. **(A)** UMAP of gene expression of 13 voltage-gated potassium channels within three FeCO clusters **(B)** Dot plot quantifying the expression levels of voltage-gated potassium channels shown in **A**. Color intensity represents mean level of gene expression in a cluster relative to the level in other clusters, and size of dots represents the percent of nuclei in which gene expression was detected.

**Fig. S5.**
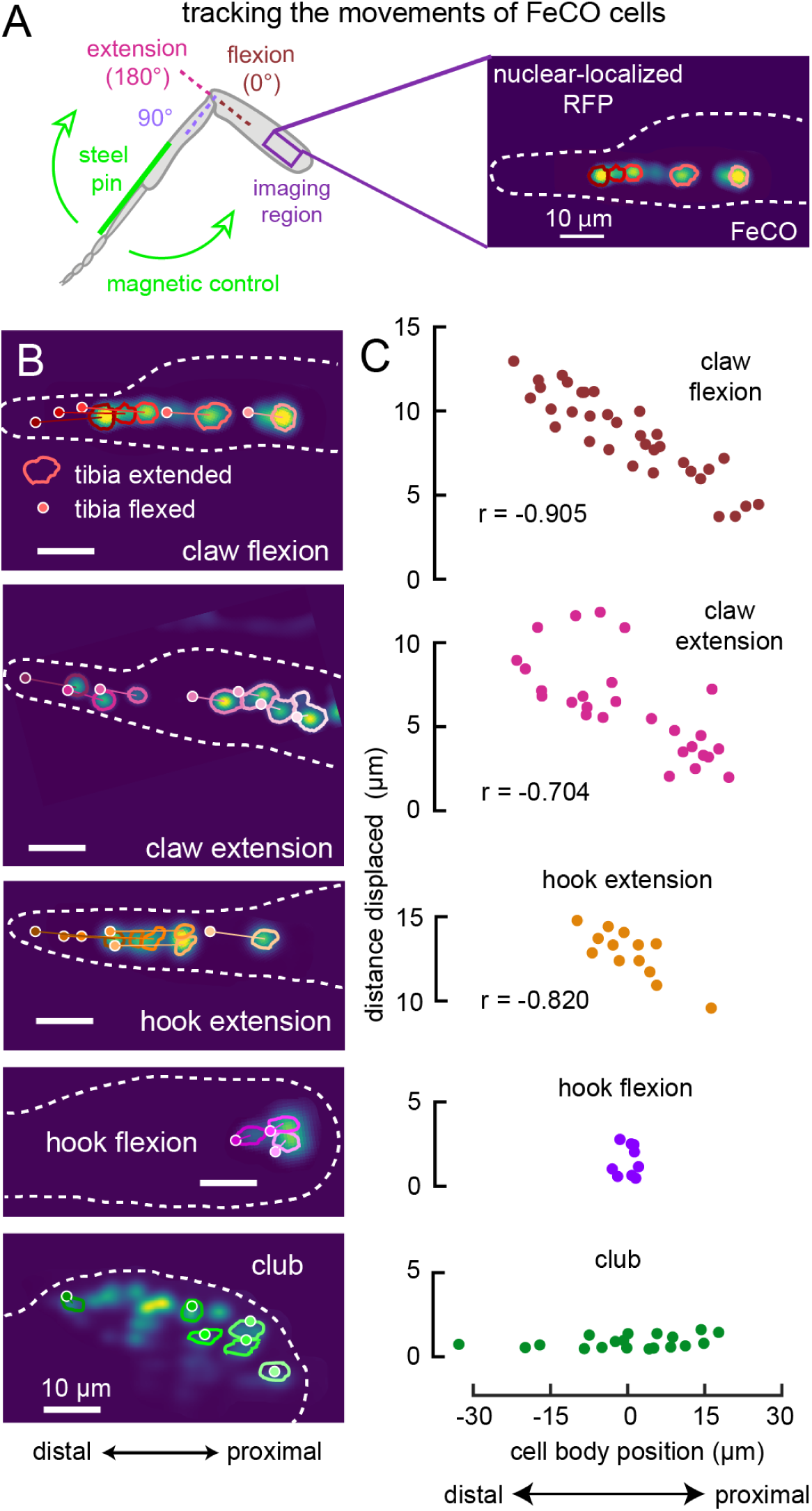
Translation of FeCO cells during movement of the femur-tibia joint. **(A)** We tracked FeCO cell nuclei labeled with nuclear-localized RFP (each colored circle in the right image indicates the tracked cell nuclei) in the femur using transcuticular two-photon imaging, while swinging the tibia from extension to flexion with a magnetic control system. (**B**) Two-photon images of FeCO cell nuclei during full tibia extension. Tracked nuclei are outlined in the images. Lines connect the centroids of each nucleus during extension (proximal end of the line) and flexion (small circles with white edge). (**C**) Scatter plots showing the distance each cell moved between flexion and extension vs. the cell’s position along the distal to proximal axis of the femur. We normalized the cell position within each fly by taking the centroid of all cells in that fly as zero and defining the distance to the proximal side as positive. (Number of flies: claw flexion = 4, claw extension = 7, hook extension = 3, hook flexion = 3, club = 3.)

**Fig. S6.**
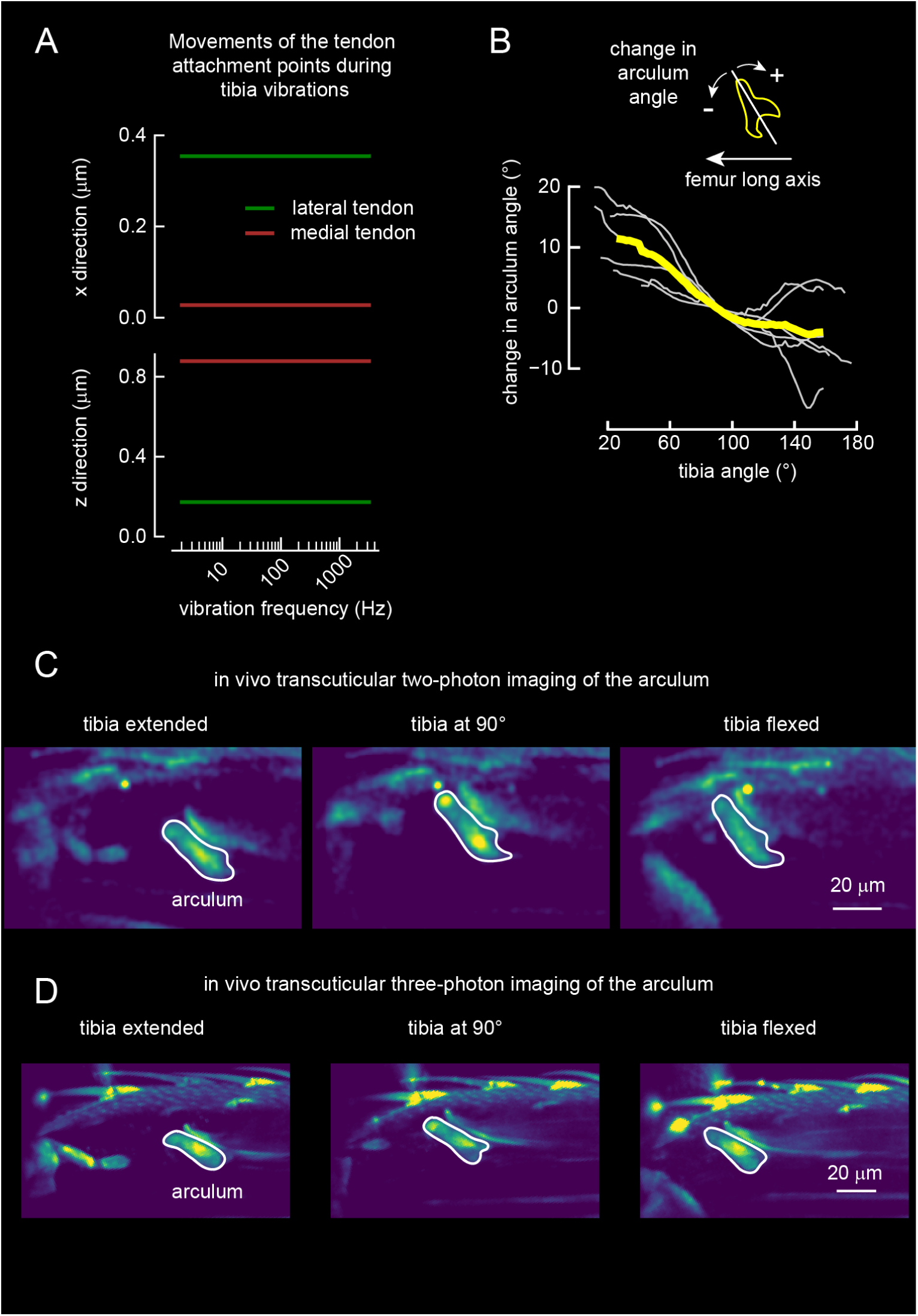
Movements of the arculum during tibia vibration and flexion/extension. **(A)** Plots showing how the attachment point for the lateral (green line) and the medial (brown line) tendon moves in the x (top) and z (bottom) direction in a 3D finite element model of the arculum receiving small periodic forces at different frequencies (1-3000 Hz). (**B**) A plot showing the change in arculum angle during full tibia flexion. The change in arculum angle is measured as the difference from the arculum angle when tibia angle is at 90°, and clockwise rotation is defined as positive change in the angle. (**C**) *in vivo* transcuticular two-photon images of the arculum at different tibia angles. White outline indicates the location of the arculum. (**D**) *in vivo* transcuticular three-photon images of the arculum at different tibia angles. White outline indicates the location of the arculum.

**Fig S7.**
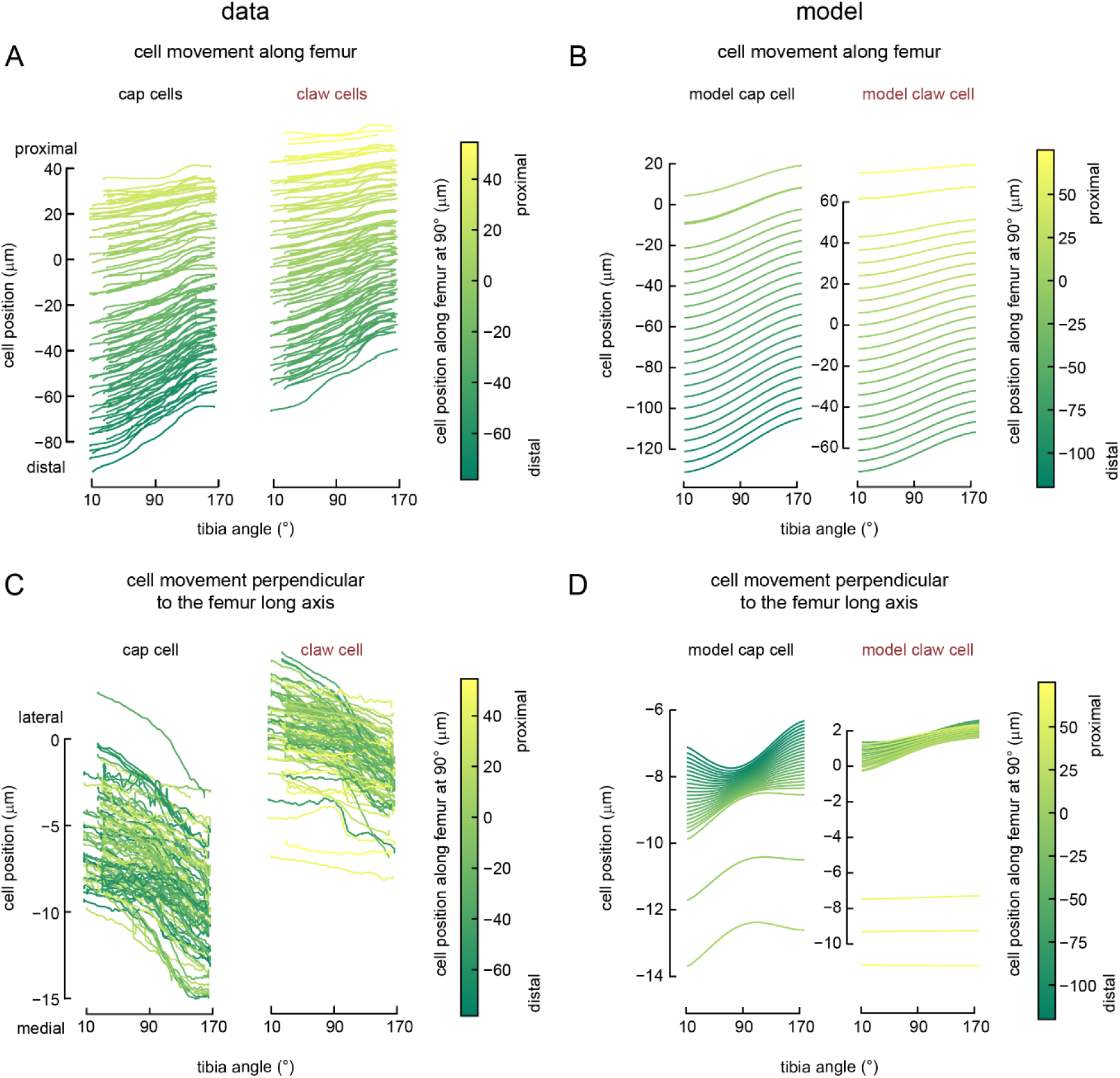
Further quantification of movements of the claw and cap cells during tibia flexion/extension for experimental and model data (same dataset and model as shown in Fig. 4). (**A**) Same as Fig. 4C. Shown here to facilitate the comparison of experimental and model data. Plots the position of cap cells (left) and claw cells (right) along the distal-proximal axis of the femur as the tibia moves from full extension to full flexion. (**B**) Same as **A,** but for the model claw and cap cells. (**C**) Same as **A**, but for movements along the axis perpendicular to the long axis of the femur. **(D)** Same as **C**, but for the model claw and cap cells.

**Fig S8.**
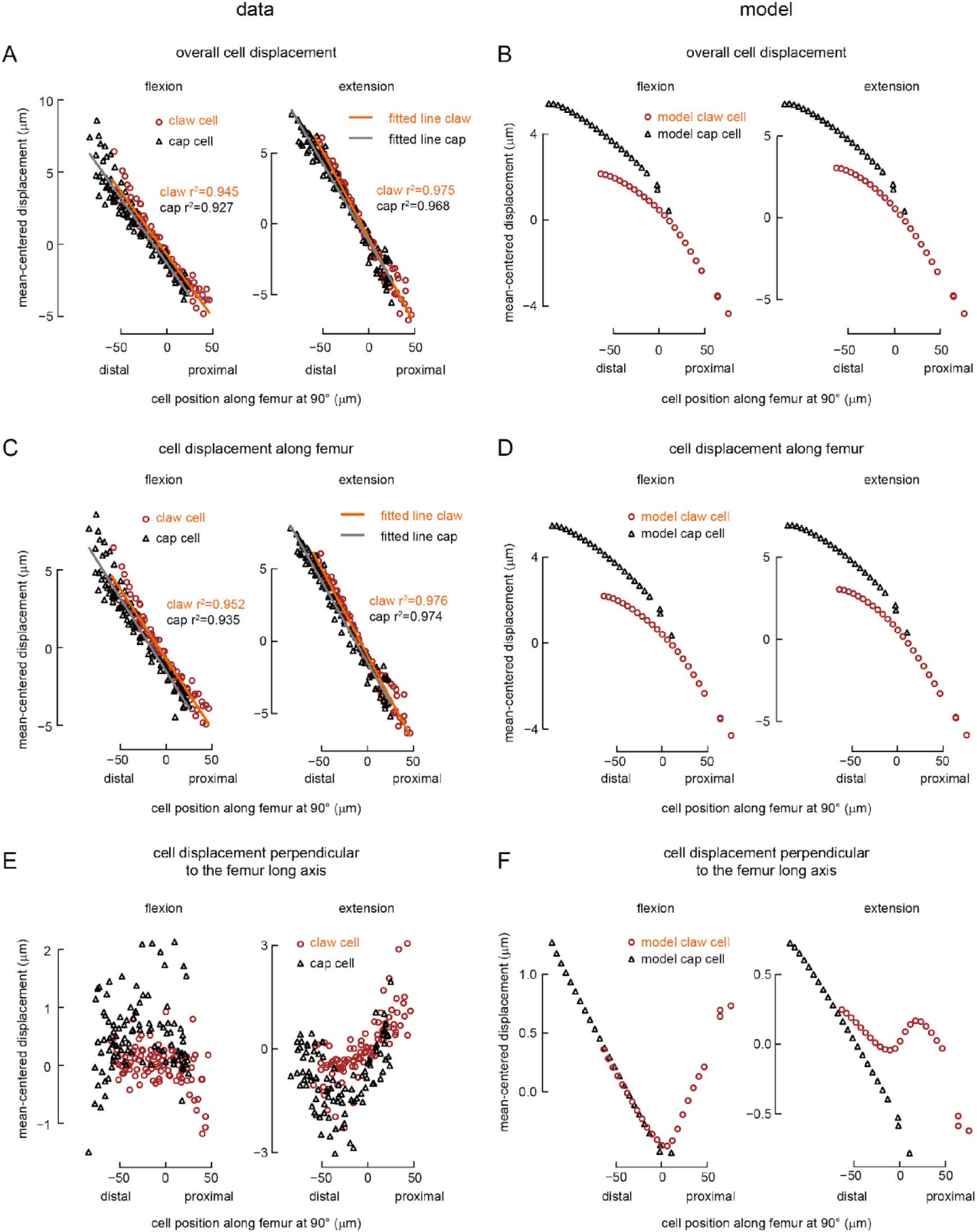
Further quantification of the displacements of the claw and cap cells during tibia flexion/extension for experimental and model data (same dataset and model as shown in Fig. 4). (**A**) Same as Fig. 4E. Shown here to facilitate the comparison of experimental and model data. Plots the overall displacement distance of claw and cap cells during tibia flexion (left) and extension (right) vs the cell’s position along the distal to proximal axis of the femur. (**B**) Same as **A**, but for the model claw and cap cells. (**C**) Same as **A**, but for the displacement distance along the long axis of the femur. For flexion, positive values indicate larger than mean displacements towards distal side. For extension, positive values indicate larger than mean displacements towards proximal side. **(D)** Same as **C**, but for the model claw and cap cells. **(E)** Same as **A**, but for the displacement distance along the axis perpendicular to the long axis of the femur. Positive values indicate larger than mean upwards displacement. **(F)** Same as **E**, but for the model claw and cap cells

**Fig S9.**
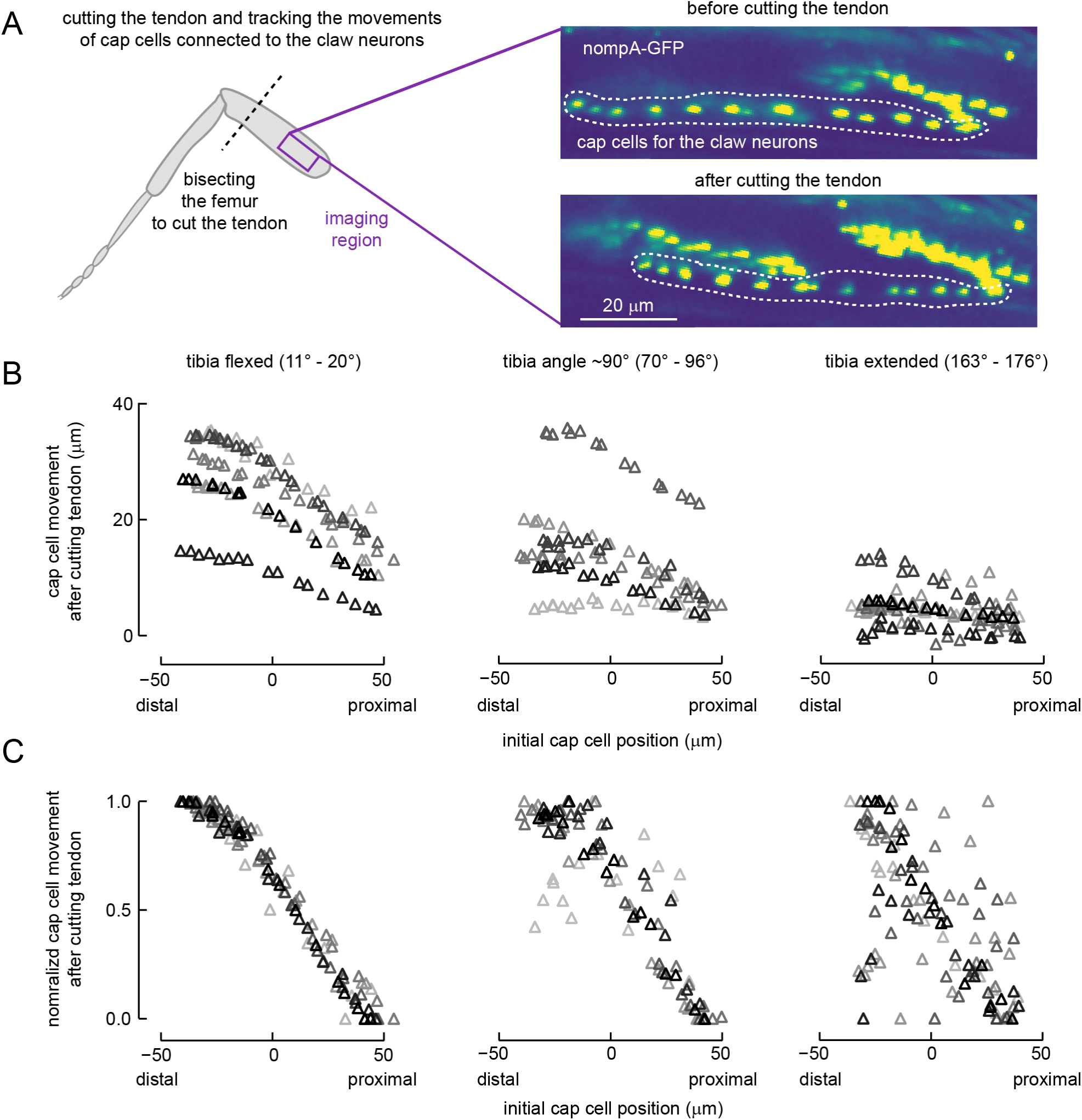
Quantification of movements of the cap cells connected to the claw neurons after manually cutting the tendon. (**A**) We glued down the femur and the tibia of a fly front leg while holding tibia at different angles. Then, we imaged the position of the cap cells connected to claw neurons (labeled with nompA-GFP) using transcuticular two-photon imaging. We cut the tendon manually by bisecting the distal femur and repeated the two-photon imaging to measure how the cap cells moved after the tendon cutting. (**B**) Scatter plots showing the distance each cap cell moved along the distal to proximal axis of the femur after the tendon cutting vs. the cell’s position along the distal to proximal axis of the femur. **Left**: Cell movements when the tibia was held at a flexed angle (11° - 20°). **Middle**: Cell movements when the tibia was held near 90° (70° - 96°). **Right**: Cell movements when the tibia was held at an extended angle (163° - 176°). Triangles with different shades of grey indicate cap cells from different legs. (Number of legs: tibia flexed = 7, tibia angle ∼90° = 6, tibia extended = 7.) (**C**)Same as A, but we normalized the cell movements within each leg.

**Figure S10.**
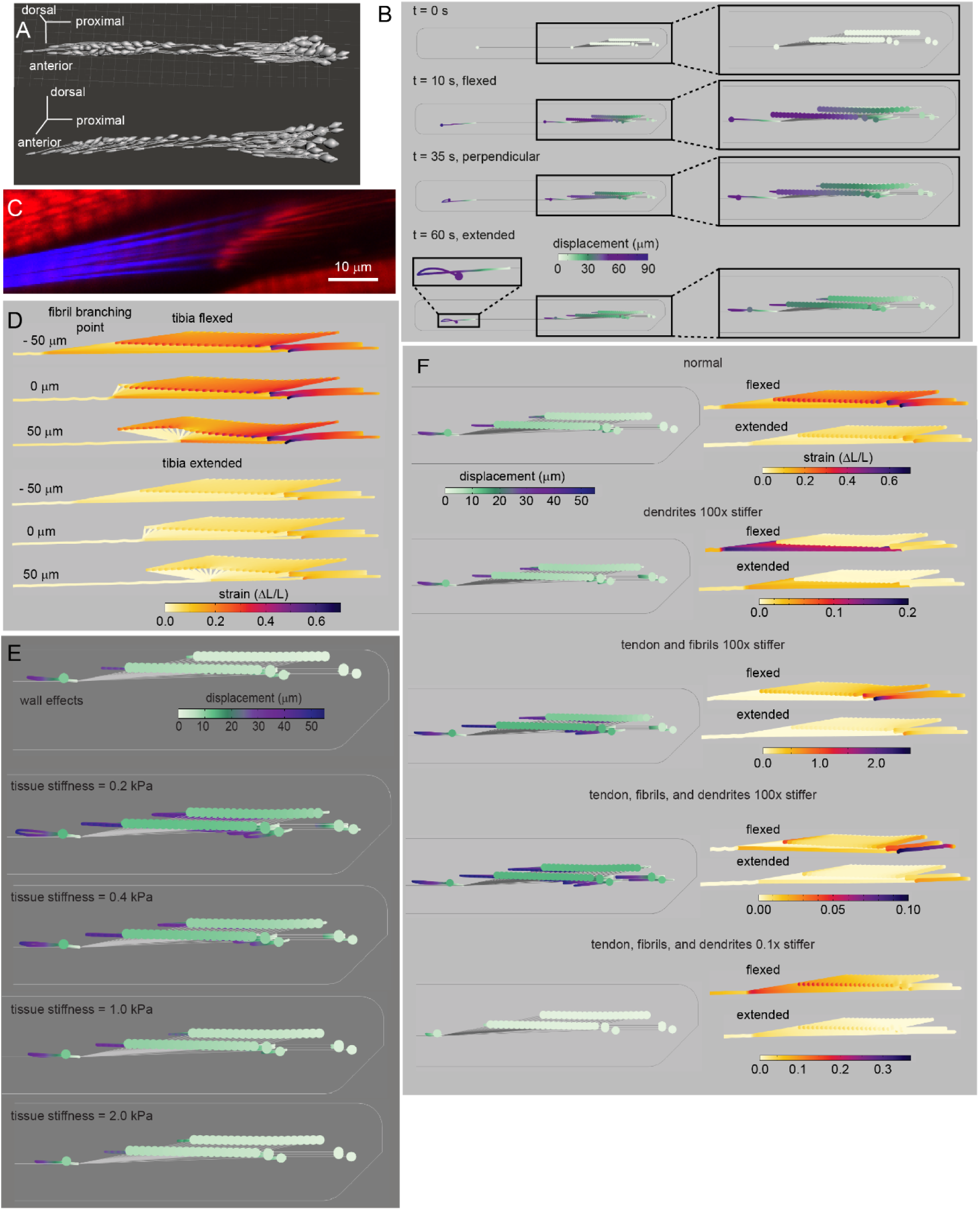
Details of the finite element model of claw neurons and tendon attachments. **(A)** A Blender model of claw and cap cells based on the X-ray reconstruction. (**B**) Cell displacement maps showing the time course of the force application. **(C)** Section of a confocal image of medial tendon fibrils fanning out (blue, autofluorescence) to attach to claw cap cells (red, phalloidin stain). **(D)** A map of strain in the dendrites of model claw cells and the fibrils connecting the dendrites to the tendon during tibia flexion (top) and extension (bottom) when we moved the fibril branching point to different positions along the long axis of the femur. When we put the branching point 50 μm proximal to the most distal cap cells, strain gradient reverses for the cells distal to the branching point. **(E)** Cell displacement maps showing how the properties of a tissue surrounding FeCO cells affect the movements of the claw cells and the cap cells during tibia flexion/extension. Having a wall closer to the claw cells reduce the movements of the claw cells. Stiffer tissues reduce the movements of all cells. (**F**) Cell displacement maps (left) and strain distribution maps (right) showing the effects of tendon and dendrite stiffness on the movements of the cells and the strains. Displacements of the cells increases when the tendon and dendrites are stiffer. Strain increases from proximal to distal dendrites as long as the stiffness of either the tendon or dendrite is normal or stiffer. If the stiffness of the both the tendon and the dendrite is lowered sufficiently, strain gradient is reversed. In all cases, strains were larger during the tibia flexion compared to during the tibia extension.

**Fig S11.**
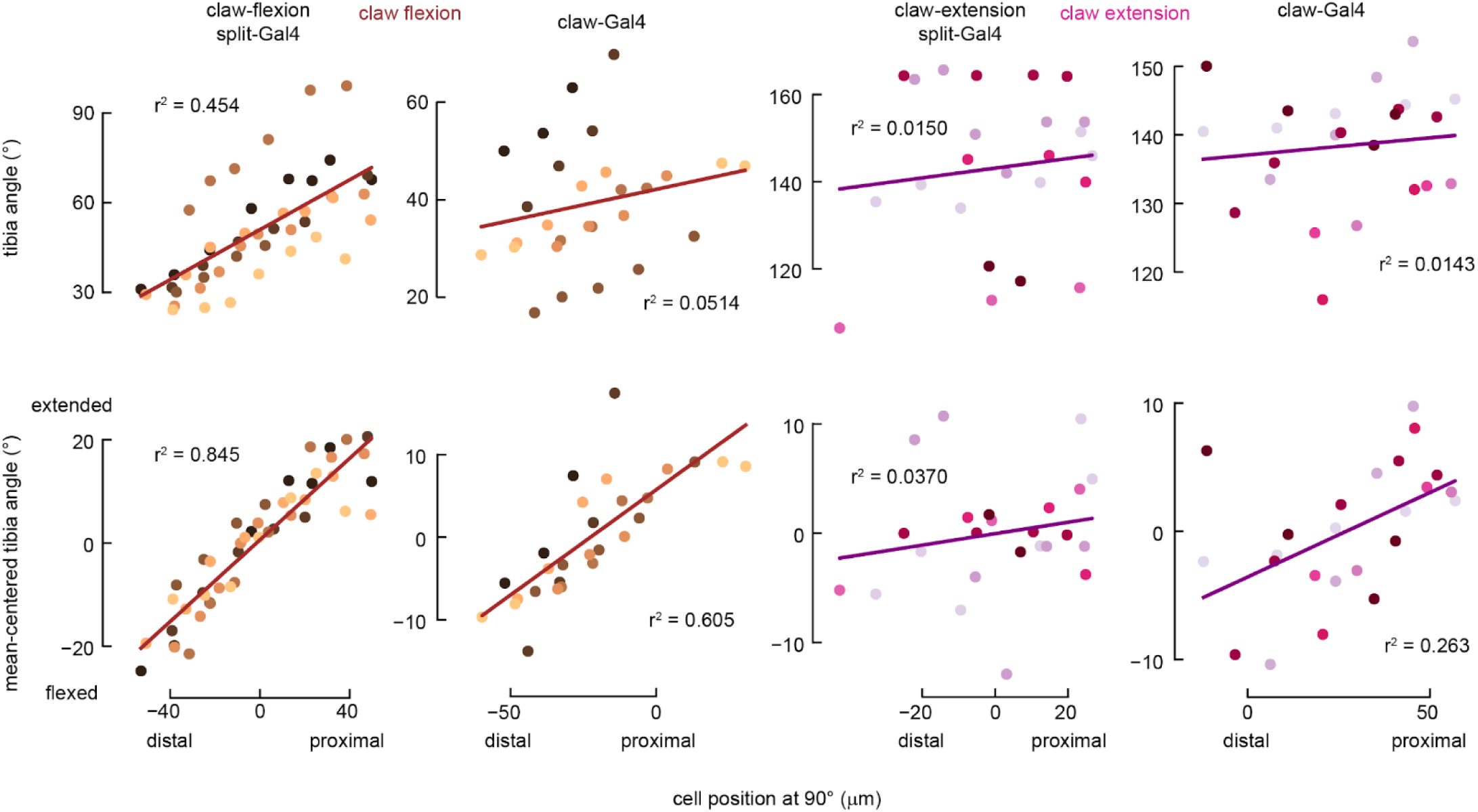
Further quantification of the relationship between the claw cells’ positions and tuning to tibia angle (**Fig. 5**). **Top row:** Scatter plots showing the relationship between each cell’s position along the proximal-distal axis of the femur and the tibia angle when the cell reached 50% of its maximum activity during a slow (6 °/s) tibia flexion (**left two columns**) or extension (**right two columns**). We labeled flexion selective claw neurons (**1^st^ and 2^nd^ column**) with two different driver lines, claw-flexion split- Gal4 line (**1^st^ column**, 7 flies; the same data as shown in **Fig. 5C**) or claw-Gal4 line (GMR73D10-Gal4; **2^nd^** column, 7 flies). We labeled extension selective claw neurons (**3^rd^ and 4^th^ column**) with claw-extension split-Gal4 line (**3^rd^ column**, 6 flies) or claw- Gal4 line (GMR73D10-Gal4; **4^th^ column**, 7 flies). Circles with different colors represent claw cells from different flies. The lines represent linear regression line. **Bottom row:** Same data as shown on the **top row**, but we mean-centered the tibia angle where the cells reached their 50% maximum activity by defining the average of this angle for all cells in each fly to be zero

**Figure S12.**
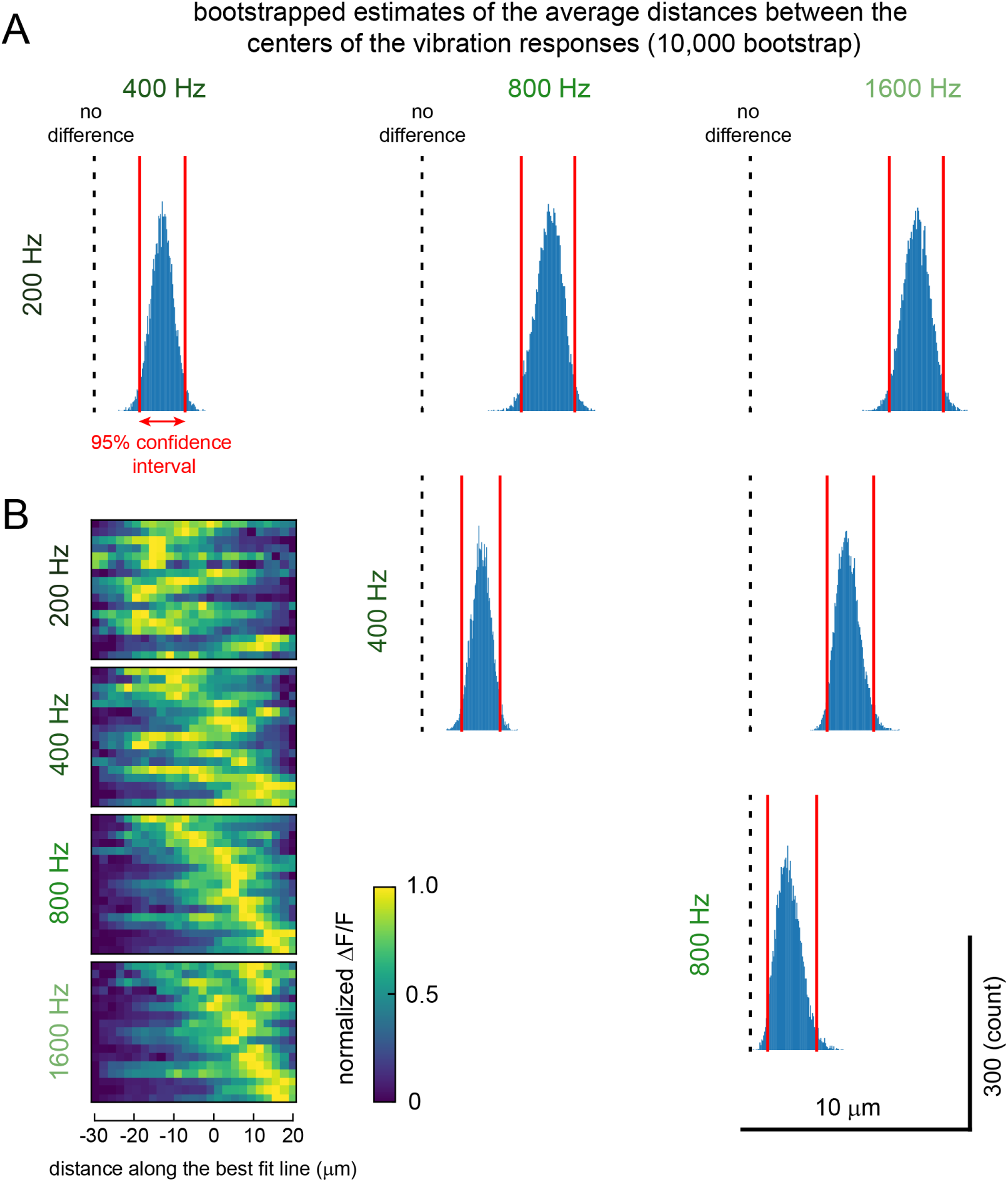
Further quantification of vibration responses in club dendrites (**Fig. 6**). Plots show bootstrapped estimates of the average distances between the centers of the responses to tibia vibrations at different frequencies and the distribution of the calcium activity along the best-fit line for the vibration response centers. **(A)** Histograms of the average distances between the centers of the responses to the tibia vibrations at different frequencies estimated with bootstrap analysis (n = 10,000 re-sampling from 17 flies). A pair of red vertical lines in each histogram show the 95% confidence interval of the average distances between the centers of the responses. Black dotted lines indicate no difference in the average location of the centers of the responses. Centers of the responses for tibia vibrations with different frequencies are all significantly different from each other. **(B)** Distribution of calcium activity (ΔF/F) along the line that best-fit the centers of the responses to tibia vibrations (a gray line in **Fig. 6D)** (n = 17 flies for each frequency). We normalized the calcium activity within each tibia vibration frequency for each fly to show the shift in the peak activity location.

**Figure S13.**
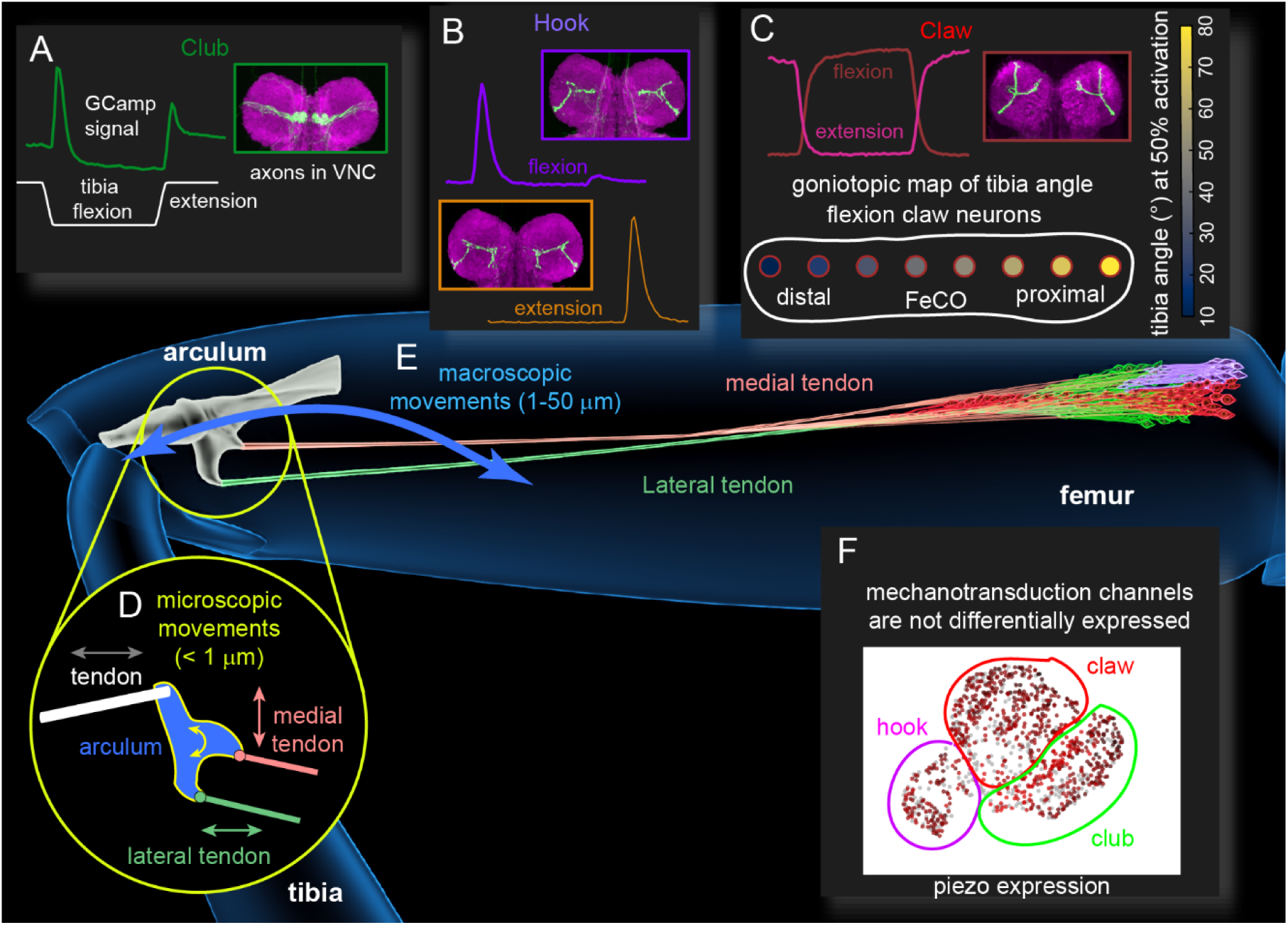
A summary of biomechanical mechanisms enabling club neurons to respond to small vibrations of the joint and claw neurons to generate a goniotopic map of tibia angle. (**A)** Club neurons are connected to the joint through the lateral tendon, and respond to small tibia vibrations and bi-directional movements of the tibia. (**B**) Hook neurons are connected to the joint through the medial tendon and respond to directional movements. Extension selective claw neurons are located towards the distal edge of the FeCO and flexion selective claw neurons are located towards the proximal edge of the FeCO. The medial tendon splits into two branches. One branch connects to the extension selective hook neurons and claw neurons, while the other branch connect to the flexion selective hook neurons. (**C)** Claw neurons are connected to the joint through the medial tendon and each claw neuron encodes specific joint angles. (**D)** A biomechanical model of the arculum suggests that during microscopic movements (< 1 μm) of the tibia, the arculum converts linear motion into a rotation. This rotation results in moving the lateral tendon parallel to the club dendrites, exciting these neurons. During the rotation, the medial tendon that connects to claw neurons moves in a perpendicular direction that does not excite the claw neurons. **(E)** During macroscopic movements (1-50 μm) of the tibia, the arculum translates along the distal-proximal axis of the femur. Medial and lateral tendons are under constant stretch tension and this tension increases as the arculum translates distally during tibia flexion. A biomechanical model of the claw neurons and their connections to the joint suggests that the geometry and the material properties of the FeCO generate larger strain on the dendrites of proximal claw neurons compared to those of distal claw neurons. This gradient in the dendritic strain leads to a goniotopic map of tibia angle represented by the activities of claw neurons as shown in **C**. (**F**) RNA-seq data suggest that different types of FeCO neurons express similar mechanotransduction channels and voltage gated sodium/potassium channels.

**Supplemental Table 1.** Top 10 genes enriched in RNA-seq clusters shown in Figure 2B

**Epithelial cell cluster**
| Gene | P-value | P-value adjusted | Log fold changes |
| --- | --- | --- | --- |
| Eip93F | 9.12E-155 | 9.40E-151 | 4.149339 |
| CG17839 | 6.28E-122 | 2.16E-118 | 2.6883016 |
| ed | 3.36E-114 | 8.64E-111 | 3.353282 |
| CG9932 | 1.80E-111 | 3.71E-108 | 3.2115743 |
| lncRNA:CR46006 | 9.67E-110 | 1.66E-106 | 3.9251654 |
| CG11147 | 2.56E-94 | 1.88E-91 | 6.127182 |
| grh | 5.90E-93 | 3.80E-90 | 6.6073465 |
| CG8312 | 6.02E-87 | 3.26E-84 | 4.916458 |
| vvl | 9.32E-87 | 4.80E-84 | 4.9913116 |
| how | 1.10E-80 | 4.06E-78 | 3.3861852 |

**Club neuron cluster**
| Gene | P-value | P-value adjusted | Log fold changes |
| --- | --- | --- | --- |
| nrv3 | 3.13E-126 | 3.23E-122 | 2.0728602 |
| Rdl | 1.16E-121 | 5.99E-118 | 2.2047484 |
| Dhc93AB | 3.73E-115 | 1.28E-111 | 3.2151954 |
| para | 2.63E-103 | 6.77E-100 | 2.9947705 |
| Sh | 1.94E-99 | 4.01E-96 | 1.901039 |
| CG12344 | 7.92E-97 | 1.36E-93 | 2.7427142 |
| CG6329 | 2.13E-92 | 3.13E-89 | 1.7301774 |
| stai | 4.46E-86 | 5.75E-83 | 2.0032818 |
| MESK2 | 1.20E-83 | 1.38E-80 | 1.0200758 |
| eag | 5.08E-77 | 5.23E-74 | 2.1537943 |

**Glial cell cluster**
| Gene | P-value | P-value adjusted | Log fold changes |
| --- | --- | --- | --- |
| N | 2.36E-48 | 2.43E-44 | 5.272594 |
| Ncc69 | 3.23E-34 | 6.65E-31 | 7.043484 |
| Pdk | 6.36E-30 | 4.10E-27 | 3.6588008 |
| CG9932 | 3.85E-29 | 2.09E-26 | 2.6037135 |
| CLIP-190 | 5.00E-29 | 2.57E-26 | 2.142356 |
| CG14687 | 7.71E-28 | 3.31E-25 | 4.3360457 |
| mtgo | 8.22E-28 | 3.39E-25 | 1.8132184 |
| Eip93F | 4.84E-27 | 1.85E-24 | 2.7748313 |
| how | 9.16E-27 | 3.37E-24 | 2.9693222 |
| IP3K2 | 2.25E-26 | 7.02E-24 | 4.4870744 |

**Claw neuron cluster**
| Gene | P-value | P-value adjusted | Log fold changes |
| --- | --- | --- | --- |
| disco-r | 5.24E-156 | 5.40E-152 | 4.6345725 |
| CG15628 | 1.39E-153 | 7.17E-150 | 3.7307715 |
| CG34384 | 1.17E-132 | 4.02E-129 | 3.3725235 |
| amon | 7.59E-132 | 1.96E-128 | 3.0669458 |
| CG43689 | 1.02E-123 | 2.10E-120 | 3.7810228 |
| Cow | 1.28E-119 | 2.20E-116 | 2.4815137 |
| CG15312 | 1.72E-114 | 2.52E-111 | 2.7650905 |
| NaCP60E | 5.89E-109 | 7.58E-106 | 2.0959513 |
| CG7236 | 5.56E-105 | 6.36E-102 | 2.412252 |
| para | 8.80E-105 | 9.06E-102 | 3.0975273 |

**Hemocyte cell cluster**
| Gene | P-value | P-value adjusted | Log fold changes |
| --- | --- | --- | --- |
| zfh1 | 8.36E-13 | 8.61E-09 | 7.131954 |
| LpR2 | 3.57E-12 | 1.84E-08 | 5.6185894 |
| Ten-m | 8.45E-12 | 2.90E-08 | 3.4689724 |
| Ppn | 1.04E-10 | 2.68E-07 | 8.746671 |
| Col4a1 | 1.35E-10 | 2.78E-07 | 7.87586 |
| fog | 3.16E-10 | 5.43E-07 | 5.115107 |
| Btk29A | 4.85E-10 | 6.27E-07 | 6.2250705 |
| pnt | 4.87E-10 | 6.27E-07 | 5.181879 |
| CG17574 | 6.25E-10 | 7.15E-07 | 5.5182843 |
| lectin-28C | 1.53E-09 | 1.57E-06 | 8.440469 |

**Hook neuron cluster**
| Gene | P-value | P-value adjusted | Log fold changes |
| --- | --- | --- | --- |
| lncRNA:CR45223 | 1.48E-69 | 1.53E-65 | 6.781883 |
| nAChRalpha5 | 1.03E-56 | 5.31E-53 | 4.6783695 |
| bru3 | 2.43E-54 | 8.35E-51 | 3.9813995 |
| Nlg3 | 8.60E-53 | 2.22E-49 | 2.7970052 |
| Ca-alpha1T | 8.40E-52 | 1.73E-48 | 4.9103622 |
| CG3339 | 1.13E-49 | 1.95E-46 | 4.082244 |
| Ptp99A | 1.95E-49 | 2.87E-46 | 1.9916081 |
| grn | 3.21E-49 | 4.13E-46 | 2.7019393 |
| jeb | 5.47E-47 | 6.26E-44 | 3.1888735 |
| Octalpha2R | 7.46E-42 | 7.68E-39 | 3.54543 |

**Muscle cell cluster**
| Gene | P-value | P-value adjusted | Log fold changes |
| --- | --- | --- | --- |
| bt | 2.31E-178 | 2.38E-174 | 8.483048 |
| up | 2.99E-177 | 1.35E-173 | 7.7078114 |
| Zasp66 | 3.94E-177 | 1.35E-173 | 7.47155 |
| sls | 7.34E-176 | 1.89E-172 | 7.646792 |
| Mf | 8.11E-174 | 1.67E-170 | 6.2663803 |
| CG33521 | 6.91E-173 | 1.19E-169 | 7.7250967 |
| CG1674 | 7.69E-172 | 1.13E-168 | 8.684115 |
| SERCA | 5.46E-171 | 7.03E-168 | 6.649068 |
| wupA | 3.22E-168 | 3.68E-165 | 8.063969 |
| Zasp52 | 4.18E-165 | 4.30E-162 | 7.40276 |

**Other neuron cluster**
| Gene | P-value | P-value adjusted | Log fold changes |
| --- | --- | --- | --- |
| erm | 2.62E-182 | 2.70E-178 | 6.1790867 |
| sosie | 2.62E-175 | 1.35E-171 | 3.685657 |
| yuri | 9.85E-162 | 3.38E-158 | 5.493425 |
| Dop1R2 | 1.75E-145 | 3.60E-142 | 4.06952 |
| CG11317 | 7.17E-143 | 1.23E-139 | 3.2864954 |
| foi | 6.54E-140 | 9.63E-137 | 2.2456117 |
| Prps | 7.32E-134 | 9.43E-131 | 2.2369118 |
| nolo | 4.69E-132 | 5.37E-129 | 2.7543526 |
| CG12065 | 3.30E-129 | 3.40E-126 | 3.3390691 |
| CG14342 | 9.04E-129 | 8.46E-126 | 3.5578682 |

## Supplemental Movies

Movie S1. Reconstruction of the sensory neurons and attachment structures of the femoral chordotonal organ (FeCO), the largest proprioceptive organ in the *Drosophila* leg.

Movie S2. Reconstruction of femoral chordotonal organ (FeCO) sensory neurons and tendons from X-ray holographic nanotomography (XNH) of the *Drosophila* leg.

Movie S3. 2-photon calcium imaging from femoral chordotonal organ (FeCO) axons in the fly ventral nerve cord.

Movie S4. Reconstruction of the arculum and femoral chordotonal organ (FeCO) tendons from X-ray holographic nanotomography (XNH) of the *Drosophila* leg.

Movie S5. Bright-field and 2-photon imaging of the arculum movements in the fly femur. Movie S6. Distribution of mechanical strain in a finite element model of claw neurons.

Movie S7. 2-photon calcium imaging from femoral chordotonal organ (FeCO) cell bodies in the fly femur.

## Materials and Methods

### Experimental animals

We used *Drosophila melanogaster* raised on standard cornmeal and molasses medium kept at 25 C in a 14:10 hour light:dark cycle. We used female flies 1 to 7 days post-eclosion for all imaging experiments, except for joint arculum imaging experiments where we occasionally used flies less than 1 day old to maximize transparency of the cuticle. The genotypes used for each experiment are listed in a table below.

### Sample preparation for confocal imaging of brains and VNCs

For confocal imaging of VNCs, we crossed flies carrying the Gal4 driver to flies carrying pJFRC7-20XUAS-IVS- mCD8::GFP and dissected the VNC from female adults in PBS. We fixed the VNC in a 4% paraformaldehyde PBS solution for 20 min and then rinsed the VNC in PBS three times. We next put the VNC in blocking solution (5% normal goat serum in PBS with 0.2% Triton-X) for 20 min, then incubated it with a solution of primary antibody (chicken anti-GFP antibody 1:50; anti-brp mouse for nc82 neuropil staining; 1:50) in blocking solution for 24 hours at room temperature. At the end of the first incubation, we washed the VNC with PBS with 0.2% Triton-X (PBST) three times, then incubated the VNC in a solution of secondary antibody (anti-chicken-Alexa 488 1:250; anti-mouse-Alexa 633 1:250) dissolved in blocking solution for 24 hours at room temperature. Finally, we washed the VNC in PBST three times, once in PBS, and then mounted it on a slide with Vectashield (Vector Laboratories). We acquired z-stacks of each VNC on a confocal microscope (Zeiss 510).

For confocal imaging of ChaT co-labeled legs, we crossed flies carrying the Gal4 driver to flies carrying UAS-RedStinger; LexAopnlsGFP/CyO; ChAT-LexA/TM6B. We selected non-balancer female adults, and dissected legs while flies were anesthetized with CO2. We immediately fixed the legs in 4% formaldehyde in PBS with 0.2% Triton-X for 20 min and rinsed them in PBS three times over 30 minutes. We mounted the legs in VectaShield and acquired z-stacks on a confocal microscope (Zeiss 510).

For *in silico* overlay of the expression patterns of Gal4 lines (**Fig. S1**), we used confocal stacks of each Gal4 line with neuropil counterstaining (from the Janelia FlyLight database (Jenett et al., 2012)) and used the neuropil staining to align the expression pattern in the VNC using the Computational Morphometry Toolkit (CMTK (Jefferis et al., 2007); http://nitrc.org/projects/cmtk) to a female VNC template (Bogovic et al., 2020).

### Sample preparation for confocal imaging of muscles, cuticle, and FeCO

To visualize the foreleg muscles, tendons and exoskeleton along with the FeCO, we expressed mCD8::GFP under control of *iav*-Gal4. Flies were anesthetized on ice, briefly washed with 70% ethanol, rinsed in PBS, and pinned onto Sylgar-coated Petri dish with 1 cm pins (Minuten Pins, Fine Science Tools # 26002-15) with their foreleg tibia in either fully levated, depressed or in an in-between position, under 2% paraformaldehyde/PBS/0,1% triton X-100, and fixed in this solution overnight at 4°C. The legs were removed at the coxa and after washing in PBS containing 1% triton X-100 (PBS-T), the samples were embedded in 7% agarose and sectioned on Leica Vibratome (VT1000s) in the plane perpendicular to the femur-tibia joint at 0.15 mm. The slices were incubated in PBS with 1% triton X-100, 0.5% DMSO, escin (0.05 mg/ml, Sigma-Aldrich, E1378), 3% normal goat serum, Texas Red-X Phalloidin (1:50, Life Technologies #T7471), anti-GFP rabbit polyclonal antibodies (1:1000, Thermo Fisher, #A10262) and a chitin-binding dye Calcofluor White (0.1 mg/ml, Sigma-Aldrich #F3543-1G) at room temperature with agitation for 24 h. After a series of washes in PBS-T the sections were incubated for 24 h in the above buffer containing secondary antibodies (1:1000, Alexa 488-cojugated goat anti-rabbit, Thermo Fisher #A32731). The sections were washed and mounted in Tris-buffered (50 mM, pH 8.0) 80% glycerol, 1% DMSO between two #1 coverslips.

### Sample preparation and confocal imaging of arculum and tendons

The flies were fixed as described above. The forelegs were removed by cutting at the trochanter-femur joint and the tibia was cut in half to create openings aiding penetration of the proteases. The soft tissues were digested away with a mixture of 0.25 mg/ml collagenase/dispase (Roche #10269638001) and 0.25 mg/ml hyaluronidase (Sigma Aldrich #H3884-100MG) in PBS-T overnight at 37°C. The cuticle’s pigmentation was bleached with 20% peroxide for 4-5 hours and the exoskeleton and tendons were stained with Congo Red (0.5 mg/mL, Sigma-Aldrich #C676-25G) overnight. The samples were dehydrated in ethanol and mounted in methyl salicylate (Sigma-Aldrich #M6752).

Serial optical sections were obtained at 1 µm intervals on a Zeiss 880 confocal microscope with a LD-LCI 25x/0.8 NA objective, or at 0.5 µm with a Plan-Apochromat 40x/0.8 NA objective. Calcofluor White, anti-GFP/anti-rabbit Alexa 488 antibodies and Texas Red phalloidin-treated samples were imaged using 405, 488 and 594 nm lasers, respectively. The 560 nm laser line was used to excite Congo Red. Images were processed in Fiji (http://fiji.sc/), Icy (http://icy.bioimageanalysis.org/) and Photoshop (Adobe Systems Inc.). Fiji was used to generate .obj files (meshes) that were further processed and segmented in Blender (http://blender.org).

### Segmentation of X-ray microscopy data

We used a previously published X-ray holographic nanotomography dataset of an adult fly’s front leg (Kuan et al., 2020). From this dataset, we manually reconstructed FeCO structures using CATMAID (Saalfeld et al., 2009; Schneider-Mizell et al., 2016); the reconstruction can be accessed at https://radagast.hms.harvard.edu/catmaidvnc/61/links/Mamiya2022. Annotated structures, including FeCO cell bodies and cap cells, tendons, and the arculum, were then exported to Blender with the CATMAID-to-Blender plugin (Schlegel et al., 2016). Unlike a previous study (Shanbhag et al., 1992), we did not observe any connections between FeCO sensory structures and surrounding muscles in the femur.

### Blender model of FeCO

We combined X-ray microscopy data mentioned above and confocal images of the foreleg exoskeleton to generate a blender model of FeCO. For modeling the arculum, we extracted the 3D model/mesh of the arculum from the confocal stack of the foreleg exoskeleton imaged with a 40x, NA 1.3 objective using Fiji’s 3D viewer plugin (http://fiji.sc/). After importing the CATMAID files for the X-ray microscopy data into Blender (Community, B.O., 2018. Blender - a 3D modelling and rendering package, Stichting Blender Foundation, Amsterdam: http://www.blender.org) as described above, we fitted the arculum mesh manually to replace the original wireframe. For the neuronal cell bodies and cap cells, we slightly modified the CATMAID primitives (spheres and cylinders) to resemble the morphology of these cells in confocal microscopy images.

### Single cell RNA sequencing and analysis

We analyzed the gene expression of FeCO neurons using single nuclei RNA sequencing (Li et al., 2022). Proximal femurs from 666 flies (iav-Gal4>unc84-2xGFP; 2-3 days old) were dissected and put into 1.5ml RNAase free Eppendorf tubes, flash-frozen using liquid nitrogen, and stored at –80°C. Single-nucleus suspensions were prepared following the protocol we described previously (McLaughlin et al., 2022). Nuclei were stained by Hoechst-33342 (1:1000; >5min). Next, we collected nuclei using Fluorescence activated cell sorting (FACS). We used the BD Aria III sorter for collecting nuclei. Since the iav+ nuclei are labeled with nuclear GFP, we gated the population first by Hoechst signal and then by GFP. Nuclei were collected into a 1.5ml tube with 200ul 1x PBS with 0.5% BSA as the receiving buffer (RNase inhibitor added). In total 5,596 GFP+ nuclei were collected. Nuclei were spun down for 10 min at 1000g at 4 °C, and then re-suspended using 43.2ul 1x PBS with 0.5% BSA (RNase inhibitor added). Since the total nuclei number after sorting is lower than the 10x maximal loading number, we loaded all the collected nuclei to 10x controller without counting.

Next, we performed snRNA-seq using the 10x Genomics system with 3’ v3.1 kit with following settings. All PCR reactions were performed using the Biorad C1000 Touch Thermal cycler with 96-Deep Well Reaction Module. 13 cycles were used for cDNA amplification and 16 cycles were used for sample index PCR. As per 10x protocol, 1:10 dilutions of amplified cDNA and final libraries were evaluated on Agilent TapeStation. The final library was sent to Novogene Corporation Inc. for Illumina NovaSeq PE150 S4 partial lane sequencing with the single index configuration Read 1 28 cycles, Index 1 (i7) 8 cycles, and Read 2 91 cycles.

Sequencing reads were aligned to the *Drosophila melanogaster* genome (FlyBase r6.31) using Cell Ranger Count (version 4.0.0). Ambient RNAs were removed using the DecontX from Celda package (version 1.10.1). Potential multiplets detected by DoubletFinder (version 2.0.3) were filtered. For further analysis of the snRNA-seq data, we utilized Scanpy (version 1.9.1). To ensure high quality data, we removed nuclei with extreme UMIs (UMI < 1000 or UMI > 3000) or high levels of mitochondrial transcripts (more than 5% of total UMIs). UMI reads were log normalized and highly variable genes were selected. Variations in mitochondrial percentage and total UMIs were regressed out, and expressions of highly variable genes were scaled to unit variance. Data were dimensionally reduced using the first 20 principal components and computed for the neighborhood graph with 10 nearest neighbors. To visualize the non-linear dimensionality reduction, we used Uniform Manifold Approximation and Projection (UMAP). Nuclei were clustered using the Leiden graph-clustering method.

### Fly preparation for *in vivo* two- or three-photon imaging of FeCO axons, FeCO cell bodies, and arculum in the VNC and leg

For recording the calcium activity of FeCO axons in the VNC while controlling tibia position, we used a fly holder and preparation procedures previously described by (Mamiya et al., 2018). First, we anesthetized the fly by briefly cooling them in a plastic tube on ice, then glued the fly onto a hole in a thin, translucent plastic sheet using UV-cured glue (Bondic). We put the fly’s head through the hole, gluing it on the upper side of the fly holder. We glued the ventral side of the thorax to the hole. Abdomen and legs were placed on the bottom side of the holder. To control the femur-tibia joint angle, we glued down the femur of the right prothoracic leg to the holder and glued a small piece of insect pin (length ∼1.0 mm, 0.1 mm diameter; Living Systems Instrumentation) to the tibia and the tarsus. We glued down all other legs away from the right prothoracic leg and bent the abdomen to the left side and glued it at that position to not interfere with the movement of the tibia of the right prothoracic leg. To enhance the contrast and improve the tracking of the tibia, we painted the pin with black India Ink (Super Black, Speedball Art Products). After immersing the top side of the preparation in *Drosophila* saline (containing 103 mM NaCl, 3 mM KCl, 5 mM TES, 8 mM trehalose, 10 mM glucose, 26 mM NaHCO3, 1 mM NaH2PO4, 1.5 mM CaCl2, and 4 mM MgCl2; pH 7.1; osmolality adjusted to 270-275 mOsm), we removed the cuticle on the ventral side of the prothoracic segment of the VNC with fine forceps. We removed the digestive tract to reduce the movements of the VNC and removed fat bodies and larger trachea to improve the optical access. We performed all recordings at room temperature.

For tracking the movements of arculum and FeCO cell nuclei or recording the calcium activity of FeCO cell bodies in the leg, we used a fly holder similar to the one described above, except that it either had a glass cover slip instead of plastic at the position where the right prothoracic femur was glued, or it was made of a thin metal sheet instead of plastic and there was a femur sized slit in the holder where the femur of the right prothoracic leg was placed. These changes allowed us to image the arculum and FeCO cell nuclei or cell bodies from above, through the cuticle, while controlling the tibia position from below the holder. For these experiments, we painted the pin with white paint to enhance its contrast against the glass cover slip or dark metal sheet. For leg imaging, we placed water between the fly holder and the objective and did not remove cuticle to avoid damaging the leg and FeCO.

### *In vivo* image acquisition two- or three-photon imaging

For imaging calcium activity from the axons of FeCO neurons and for tracking FeCO cell nuclei movements, we used a modified version of a custom two-photon microscope previously described in detail (Euler et al., 2009). We used a mode-locked Ti/Sapphire laser (Mira 900-F, Coherent) set at 930 nm for the excitation source and adjusted the laser power using a set of neutral density filters to keep the power at the back aperture of the objective (40x, 0.8 NA, 2.0 mm wd; Nikon Instruments) below ∼25 mW during the experiment. For calcium imaging from the FeCO cell bodies in the leg and for simultaneous imaging of the cap cells and the claw cell nuclei, we used a Movable Objective Microscope (MOM; Sutter) with a mode-locked Ti/Sapphire laser (Coherent Vision) set at 920 nm for the excitation source and adjusted the laser power using a Pockels cell to keep the power at the back aperture of the objective (40x, 0.8 NA, 3.5 mm wd; Nikon Instruments) below ∼25 mW during the experiment. For autofluorescence imaging of the arculum in the leg, we either used the two-photon Movable Objective Microscope mentioned above, or the three-photon excitation microscope. For three-photon excitation microscope, we generated excitation laser pulses with a commercial non-collinear OPA (Pera-F, Coherent) operating at 1300 nm pumped by a 40W fiber laser (Monaco, Coherent) operating at 1040 nm. We controlled the laser scanning mirrors and image acquisition of all microscopes with ScanImage software (version 5.7) (Pologruto et al., 2003) running in MATLAB (MathWorks).

To detect GCaMP7f fluorescence and the autofluorescence from the arculum, we used an ET510/80M (Chroma Technology Corporation) emission filter (VNC imaging) or a FF03-525/50-30 (Semrock) emission filter (cell body imaging / arculum imaging). For detecting RFP or tdTomato fluorescence, we used a 630 AF50/25R (Omega optical) emission filter (VNC and cell nuclei imaging) or a FF02-641/75-30 (Semrock) emission filter (cell body imaging). We amplified the fluorescence signals with GaAsP photomultiplier tubes (H7422P-40 modified version without cooling for VNC and cell nuclei imaging; H10770PA-40 for leg imaging and the arculum imaging; Hamamatsu Photonics).

For VNC axon recordings and cell nuclei tracking, we acquired images (256 x 120 pixels) at 8.01 Hz from the axon terminals of major axon bundles of FeCO neurons, or from the proximal region of femur where FeCO cell nuclei are located. For recordings of FeCO cell bodies, dendrites, or the arculum, we acquired fast z stacks of images (for cell bodies, 512 x 250∼300 pixels, 4∼7 z-levels; for dendrites, 512 x 512 pixels, 6 z-levels; for the arculum, 512 x 256 pixels, 7 z-levels) using resonant scanner and piezo controlled objective (acquisition rate for cell bodies, 9.6∼16.8 Hz; for dendrites, 5 Hz; for the arculum, 8.51 Hz). We adjusted the image size and stack depth to capture most of the cell bodies, dendrites, or the arculum throughout each trial. At the end of the experiment, we acquired a z stack of the imaged regions to confirm the recording location.

### Controlling tibia position using a magnet-motor system

We used a magnetic control system previously described in detail (Mamiya et al 2018) to control the femur-tibia angle during both the recordings from the axon terminals in the VNC and the cell bodies in the leg. We moved the tibia/pin to different positions by magnetically pulling them using a rare earth magnet (1 cm height x 5 mm diameter column). To precisely control the speed and position, we attached the magnet to a steel post (M3 x 20 mm flat head machine screw) and controlled its position using a programmable servo motor (SilverMax QCI-X23C-1; Max speed 24,000 deg/s, Position resolution 0.045 deg; QuickSilver Controls). To vibrate the tibia at high frequency, we placed a piezoelectric crystal (PA3JEW; ThorLabs) between the magnet and the steel post. We placed the motor on a micromanipulator (MP-285, Sutter Instruments) and adjusted its position so that the magnet moved in a circular trajectory centered at the femur-tibia joint, with the top edge of the magnet at approximately the same height as the tibia/pin. The inner edge of the magnet was ∼1.5 mm from the center of the femur-tibia joint and the distance between the pin and the magnet was ∼ 300 µm. We controlled the speed and the position of the servo motor with a custom script written in QuickControl software (QuickSilver Controls). For each trial, we confirmed the movement and the position of the tibia-femur joint using a high-speed video camera and machine vision algorithms described below.

Because it is difficult to fully flex the femur-tibia joint without the magnet colliding with the abdomen or other legs, we only flexed the joint up to ∼ 18°. For swing motion trials, we commanded the motor to move between fully extended position (180°) and fully flexed position (18°) in one continuous sweep at a set speed of 360 deg/s (axon terminal imaging) or 5 deg/s (cell body imaging). For faster swing trials, we had a 5 s interval between the flexion and the extension movements, and we repeated the swing motion 6 times with a 5 s inter-trial interval. For slower swing trials, we had a 2 s interval between the flexion and the extension movements, and we repeated the swing motion 3 times with a ∼30 s inter-trial interval. Because responses to each repetition of the swing motion were similar, we averaged the responses to the swing motion across the trials.

For ramp-and-hold motion trials (axon terminal imaging), we commanded the motor to move in 18° steps between the fully extended position and fully flexed position at 240 deg/s. Between each ramp movements, the leg was held at the same position for 3 s. We repeated each type of ramp-and-hold motion 3 times and averaged the responses across trials within each fly.

For both types of movements, we set the acceleration of the motor to 72000 deg/s^2^. Movements of the tibia varied slightly across trials and flies due to several factors, including a small offset between the center of the motor rotation and the femur-tibia joint, resistance to the magnetic force by the fly, interference by physical contact between the tibia/pin and the holder or abdomen/legs. Because these variations were relatively small, we did not consider these differences in the summary of the responses (**Fig. 1D**, **S2F**). However, when quantifying the relationship between the femur-tibia angle and the cell activity (**Fig. 5C, S11**) or movement (**Fig. 4C, S7**), we plotted the response against the actual femur-tibia angle during each trial.

### Tracking the femur-tibia joint angle

To track the position of the tibia/pin during the trials, we recorded videos (200 fps) using an IR sensitive high speed video camera (Basler Ace A800-510um, Basler AG) with a 1.0x InfiniStix lens (94 mm wd, Infinity). For VNC recording experiments, we used an 850 nm IR LED (M850F2, ThorLabs) to backlight the black painted tibia/pin through the translucent plastic fly holder. For leg recording experiments that required the metal fly holder, we front-lit the white painted tibia/pin using the same LED. In both cases, we used a 900 nm short pass filter (Edmund Optics) to filter out the two-photon excitation laser light. Because the servo motor was placed directly under the fly, we placed the camera to the side and used a prism (Edmund Optics) to capture the view from below. To synchronize the camera’s images with two-photon microscope’s laser scanned images, we acquired both the camera’s exposure signal and the galvo scanner’s Y axis scanning signal at 20 kHz.

For tracking the black painted tibia/pin position in the backlighted image (VNC recordings), we first selected the pixels below the threshold to detect the tibia/pin. We then approximated the orientation of the tibia/pin as the long axis of an ellipse fitted to these pixels (Haralick and Shapiro, 1992). In the front-lit setup (cell body recordings), lighting varied significantly throughout the trial due to shadows cast by the fly’s body and the magnet. Because of this, we used DeepLabCut (Mathis et al., 2018) to track the tibia/pin. We trained the neural network to track 5 points along the femur and tibia/pin and approximated the orientation of each body part by fitting a line through the tracked 5 points using principal component analysis. The spatial resolution of the camera image was 3.85 µm per pixel and, assuming circular movement of the tibia/pin, 1-pixel movement at the edge of the tibia/pin (∼1.2 mm from the center of the rotation) corresponded to 0.18°.

### Vibrating the tibia using a piezoelectric crystal

To test the responses of the axons and dendrites of FeCO neurons to high frequency vibrations of the tibia, we attached the magnet to the tibia/pin and moved it using a piezoelectric crystal (PA3JEW, Max displacement 1.8 µm; ThorLabs). For controlling the movement of the piezoelectric crystal, we generated command waveforms in MATLAB and sent them to the piezo through a single channel open-loop piezo controller (Thorlabs). Because previous calibration of this piezoelectric crystal’s movement in response to sine waves of different frequencies (Mamiya et al., 2018) showed that the power of oscillation at the target frequency drops greatly when the frequency of the command sine wave exceeds 2000 Hz, we used vibration in the range of 100 – 1600 Hz (sampling frequency 10 kHz). For each frequency, we presented 4 s of vibration twice with an inter-stimulus interval of 8 s.

### Processing and analysis of *in vivo* imaging data

We performed image processing and analyses using scripts written in MATLAB (MathWorks) and Python. For all recordings, we first applied a Gaussian filter (kernel size 5x5 pixel, σ = 3) to the acquired images to reduce noise. For the recordings of FeCO axon terminals in the VNC, we aligned the filtered images (registered to ¼ pixel) to a mean image of the trial using a sub-pixel image registration algorithm (Guizar-Sicairos et al., 2008). For this alignment, we used the tdTomato fluorescence, which does not change as a function of intracellular calcium concentration. tdTomato fluorescence was stable over the course of the experiment, indicating that movement artifacts were absent or small for the VNC recordings. Thus, for these experiments we used the change in GCaMP7f fluorescence relative to the baseline (ΔF/F) as the indicator of the calcium activity. To calculate ΔF/F, we first selected pixels whose mean GCaMP7f fluorescence was above a set threshold (150 arbitrary units) and used the lowest average fluorescence level of these pixels in a 10-frame window as the baseline fluorescence during the trial. Because split-Gal4 lines drove GCaMP7f in one functional type of FeCO neurons, we averaged the responses of all pixels for these experiments. For claw-extension split-Gal4 line that drove expression in primarily extension-tuned claw neurons, but also occasionally labeled flexion-tuned neurons, we first separated the selected pixels into two groups based on the similarity of their intensity change during the trial using k-means clustering of pixel correlations (Mamiya et al., 2018). We then calculated the average Δ F/F for each group of pixels. When analyzing vibration responses in the axons, we took an average activity level in a 1.25 s window starting 1.25 s after the vibration onset as a measure of response amplitude. We further averaged the responses within each fly before averaging across flies.

For recordings of FeCO cell nuclei and cell bodies in the leg, we found that different cell bodies move by different amounts during the flexion/extension of the joint. Thus, we did not attempt to align these images to the mean image for these recordings. Rather, we used a cell tracking algorithm described below, or manual labeling of each cell nuclei in the image to track the position of each cell during the trial. Most of the cell movements occurred in the x-y direction of the imaging plane. Thus, for recordings with fast z-stacks (cell body imaging), we first averaged the images acquired from the different z-levels for each time point to be able to track cell bodies at the different z-levels simultaneously. Then we selected pixels whose mean RFP (cell nuclei tracking) or tdTomato (cell body imaging) fluorescence was above a set threshold (20 – 150 arbitrary unit), resulting in groups of pixels shaped similar to the cell nuclei or cell bodies with higher intensity fluorescence in the middle of the cell. We used this fluorescence intensity pattern to perform a watershed segmentation of each image to identify cell nuclei or cell bodies. We also used the same algorithm to identify the autofluorescence from the arculum in each image and track it during tibia flexion/extension. Because simple watershed segmentation often leads to over-segmentation of the image, we first chose a key frame in the trial, and manually merged the over-segmented sections so that the segmentations matched the cells in this image. Then, we used the centroids of these segmented cells as the seeds to perform watershed segmentation of the image corresponding to the next frame. Because the cell movements between each imaging frame were small compared to the cell size (for slow swing trials), this procedure allowed us to track the movements of the cell by updating the centroid of each cell as it moved. We repeated this process until we labeled all frames in this manner. To reduce noise in the labeling process, we selected three key frames per trial (located at the beginning, middle, and the end of the trial) to generate three different sets of labels for each frame. Then we merged these labels together by taking pixels whose label identity agreed in at least two of the three labels. Watershed segmentation relies on the intensity pattern of the fluorescence for segmentation, and because some lines label cells that are in close proximity, it was difficult to segment out these overlapping cells. In these cases, we chose to group these cells together. Thus, our segmentation is a conservative estimate of the cells in the recordings, and may sometimes contain segments that are composed of multiple cells. Because tdTomato fluorescence fluctuated as the cell moved inside the leg (presumably due to different parts of cuticle and leg transmitting different amounts of light), we used the changes in the ratio between the GCaMP fluorescence and tdTomato fluorescence relative to the baseline ratio (ΔR/R) from each cell as the indicator of the calcium activity. For each cell, we used the lowest average ratio in a 12-frame window as the baseline ratio during the trial. For cells labeled with the claw-extension split-Gal4 line that occasionally drove expression in flexion-tuned neurons and those labeled with the claw- Gal4 line (GMR73D10), we categorized the cells into flexion-tuned or extension-tuned based on their peak responses to tibia angle.

### Analyzing the relationship between the position and activity of claw neuron cell bodies

To investigate the relationship between the position of claw neurons and their activity patterns, we measured each segmented claw neuron’s position along the long axis of the femur. Cell bodies of the claw neurons are aligned in a blade-like array along the long axis of the femur and we approximated this axis by fitting a line through the centroid of the segmented cells using principal component analysis. We used the position of each segmented cell body’s centroid along this axis when the tibia angle was 90° (average of the cell’s position when 87.5° <= tibia angle <= 92.5°) as a measure of the cell’s position. Because claw neurons’ maximum response amplitudes varied from neuron to neuron, we looked at the relationship between the tibia angle when the claw neuron’s activity has reached 50% of the maximum activity (**Fig. 5B-C, S11**).

### Quantifying the map of vibration frequency in the dendrites of club neurons

Within the FeCO, the dendrites of the club neurons responded most strongly and consistently to tibia vibration. Thus, we focused our analysis to the club dendrites and manually identified the region that corresponds to the dendrites of the club neurons (ROI) at each z-level for each fly (6 z-levels). To quantify the spatial pattern of activity, we first measured the changes in GCaMP7f fluorescence during the tibia vibration relative to the baseline (ΔF/F) for each ROI at each z-level. Because most of the changes in the activity pattern was in x-y direction, we next took an average ΔF/F value along the z-axis to generate a 2D map of the activity pattern and calculated the weighted center of the activity. To compare the activity pattern between flies, we re-oriented the ΔF/F map so that the club dendrites were horizontal in all the maps. We also normalized the location of the center of the activity by using the center point of the dendritic tips of club neurons as the origin. To determine the center point of the dendritic tips, we looked at the most distal layer of dendritic tips and took the midpoint between the most dorsal and ventral dendritic tips.

For bootstrapped estimates of the average distances between the centers of the vibration responses, we first calculated the distances between the response centers for different pairs of vibration frequencies (from 200 to 1600 Hz) in each fly (n = 17 flies). Then, for each pair of vibration frequencies, we resampled the distances with replacement and calculated the average distance between the response centers 10,000 times. We looked at the 95% confidence intervals of the average distances between the response centers and checked whether the confidence intervals overlapped with zero distance (no difference between the locations of the response centers).

For calculating the response distribution along the best-fit line, we binned the pixels based on the distance along this line (2 µm/bin) and averaged their ΔF/F values. For each vibration frequency in each fly, we normalized the response distribution by its maximum response (n = 17 flies). In each distribution map, we ordered the flies based on the maximum response location during the 800 Hz tibia vibration.

For calculating the frequency response turning curve, we used pixels in the ROI that had the baseline fluorescence level above the background (background fluorescence level in each prep was identified by plotting the histogram of pixel intensities and finding a threshold that best distinguish background pixels from those that have GCaMP7f expression. Typically, there are two distributions of the pixel intensity, one that corresponds to the background pixels (narrow distribution at the low pixel intensity) and wider distribution of brighter pixels that corresponds to GCaMP7f expression). We calculated the average change in GCaMP7f fluorescence relative to the baseline for these pixels and used it as a measure of the activity (average for two stimuli in each fly for each frequency).

### Quantifying the movements of the cap cells after cutting the tendons

To determine the direction of the forces acting on the cells of FeCO during tibia flexion/extension, we cut the tendon manually while holding tibia at different angles and observed how the cap cells move. To immobilize the fly with tibia at different angles, we used the same fly holder as in the imaging experiments, but kept the two front legs on the upper side of the holder and glued the femur and tibia of each leg to keep the tibia to either flexed (11° - 20°), near 90° (70° - 96°), or extended (163° - 176°) angle. After gluing down the fly, we took a digital image of the preparation and visually confirmed the tibia angle. Then, we used the same two-photon imaging setup as in the VNC axon recording experiments described above to take a z-stack (512 x 512 resolution, 45 to 68 steps, 1 μm per step) of the cap cells expressing GFP (nompA-GFP flies). To cut the tendon, we covered the preparation with the fly saline and cut the entire femur at about 1/3 from the distal end using 30G hypodermic needle (BD). After cutting the tendon, we took another z-stack of the cap cells to observe the change in the cell position.

To quantify the movements of the cap cell after cutting the tendon, we manually marked the position of the cap cells in each z-stack using napari (Sofroniew, Nicholas et al., 2022). Because cutting the femur sometimes resulted in slight shifts in the position of the femur, we also marked the roots of bristles in the z-stacks and used them as a landmark to align the two z-stacks with rigid translation and rotation using least square fitting (Arun et al., 1987). Cap cells are aligned in a blade-like array along the long axis of the femur and we approximated this axis by fitting a line through the marked cap cell positions using principal component analysis. We used the cap cell’s position along this line as the measure of the cap cell position (mean-centered by subtracting the average position of the cell) and we used the movements of the cap cells along this line as the measure of the cap cell movement (measured the movement towards the proximal side as positive movements).

### A finite element model of the arculum

We conducted a finite element (FE) analysis using a series of FE models to study how the arculum moved in response to the forces applied to it *via* the femur-tibia joint. This model was implemented using COMSOL Multiphysics (ver.5.5, Massachusetts, USA). Models were developed in the structural mechanics module and all components were modeled as simple linear elastic materials with linear behavior that follows Hooke’s law.

We based arculum geometry on the 3D Blender model of the arculum and tendons reconstructed from confocal images and X-ray microscopy data (**Fig. 3**). This mesh was then imported into Meshmixer. After removing the joint and tendon attachments, we generated an STL file and imported the model into COMSOL. Based on the confocal images, we identified sections of the model that tendon attaches to and marked them accordingly.

The 3D model was meshed using tetrahedral elements with triangular surface faces. We used the extremely fine setting of the physics controlled meshing. This uses a minimum element of 0.63 um and allows a maximum element size of 3.5 um. The model uses smaller elements in narrow regions of the arculum, specifically where the medial and lateral tendons attached to it in order to more carefully calculate strains and stresses as they develop in these regions. The complete geometry was represented by >35k elements.

### Material properties in the arculum model

We modeled the arculum as a linear elastic material. Most insect cuticle lies within a narrow range of densities of ∼1200 kg/m^3^ (Vincent and Wegst, 2004) and the arculum was modelled with this density. We used Poisson’s ratio of 0.3, which is common for most materials in nature. The exact Young’s modulus of the arculum is not known. Video and UV fluorescence data suggests that it contains some resilin but is stiffer than the resilin containing tendons that anchor it within the femur. There are different estimates for the Young’s modulus for resilin and here we use 1.8 MPa (Frantsevich et al., 2019; Gosline et al., 2002). For our main model, we use a Young’s modulus of 3.6 MPa (twice that of resilin) for the arculum. We also performed a parameter sensitivity study for this input parameter and found that this value predicted the best vector decomposition. In the femur, the arculum is suspended within femoral tissue and is expected to encounter a high level of damping due to viscosity of the local tissue. Thus, to mimic these conditions, we applied a high isotropic damping factor of 0.5 to the arculum.

### Boundary conditions of the arculum model

The arculum is suspended in place in the femur by 4 tendon attachments. These are the joint tendon from the tibia-femoral joint, the extensor tendon to the extensor muscle within the femur, and the medial and lateral tendons to the FeCO. We treated each of these attachments as the boundary conditions experienced by the arculum. Each attachment is modelled as a spring and the attachment points are demarcated on the arculum model. The joint, medial, and lateral tendons are modelled as running parallel to the long axis of the femur (x axis) and the extensor tendon is modelled as making a 45° angle between the x and z axes. We treated each tendon as a stiff element whose spring constant is calculated using the following formula,

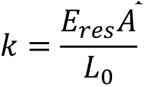

Where, Eres is the Young’s modulus of resilin, L0 is the equilibrium length of the tendon, and “*A*” is the cross sectional area of the tendon. Based on the X-ray microscopy data, we used the following values to model each tendon:

| Tendon name | Equilibrium length ($\mu\text{m}$) | Cross sectional area ($\mu\text{m}^2$) |
| --- | --- | --- |
| Joint tendon | 72 | 295.3 |
| Femoral tendon | 225 | 295.3 |
| Medial tendon | 268 | 50.9 |
| Lateral tendon | 283 | 57.2 |

We observed that the proximal edge of the medial tendon attached to the FeCO undergoes large translation movements during joint flexion (**Fig. 3F, G**, **S6**). This suggests that this tendon is effectively coupled to a soft spring on the proximal side. To model this, we treated this tendon as having a spring constant lowered by a factor of 2. We also performed a parameter sensitivity analysis to test the effect of this approximation and found that the arculum’s motion was unaffected. Finally, the spring foundations were initialized as being in equilibrium, i.e. not applying any forces on the arculum unless moved from initial positions. Each spring foundation was also modelled as experiencing significant damping along its axis (isotropic loss factor =0.5).

### Loads generated during joint rotation in the arculum model

During joint flexion, loads are applied to the arculum and these loads are in turn transmitted to the FeCO via the medial and lateral tendons. Bright-field imaging (**Movie S5**) showed that the joint tendon converts the rotation experienced by the tibia into a linear movement. This suggests that the joint tendon converts joint torque into a linear force along the long axis of the femur (x axis). We modelled this force as a periodic 10 uN force applied at the attachment area of the joint tendon parallel to the x axis. This approximates force levels known to be produced by *Drosophila* muscles (Azevedo et al., 2020). Since the model follows linear Hookean mechanics, an increased force should simply lead to a linear increase in the movements of the arculum, as long as the forces are below the point where the force cause plastic deformation or buckling of the arculum. We tested the behavior of the model for a period force from 2 Hz to 80 kHz, where the 2^nd^ eigenfrequency of the system was observed. We used a frequency domain study, simulating either 20 or 5 frequency steps per frequency decade depending on the study.

### FeCO models to investigate joint angle selectivity in claw cells

We used a series of finite element models to study the biomechanical mechanisms underlying the movements of claw neurons and the development of the dendritic strain gradient during the tibia flexion/extension. We implemented these models using COMSOL Multiphysics (ver. 6.1, Massachusetts, USA). The models investigate how the geometry of the FeCO and the stiffness of the fibrils, dendrites, and surrounding tissues give rise to the observed cell movements and the subsequent development of the strain gradient in the dendrites (**Fig. 4, S7, S8**). There are two sets of claw cells and dendrites in the FeCO, and a pair of claw cells is attached to each cap cell which also forms an array (**Fig. S10A**). Each array of claw cells lies in a plane, and the two planes are about 20 degrees apart from each other. The dissecting plane between these two cell arrays lies along the ventral dorsal axis of the femur. Thus, forces applied by the medial tendon along the dorsal ventral and proximal distal planes, would be symmetrically applied to both arrays of neurons. To reduce the size of the computational problem, we believe it is reasonable to use the symmetry between the two neuronal arrays and simplify this geometry into a 2D system. Therefore, we represent both arrays of cells by a single in planar array which maintains all other geometric features of each array, specifically the spacing of the claw cells along the proximal distal axis and the angle of the dendrites as well as the fanned array of tendon fibrils attached to this array of cap cells (**Fig. 4D**).

We developed and solved the 2D models using the truss and solid mechanics interfaces of the structural mechanics module in COMSOL Multiphysics (ver. 6.1, Massachusetts, USA). We modelled the medial tendon, the fibrils, and the claw cell dendrites as truss elements, i.e. as slender ‘string or cable’-like structures with reasonable axial stiffness, but low to negligible bending stiffness. Such behavior would most closely resemble the behavior of soft structures like tendons or neuronal dendritic cilia. We modelled the tissue surrounding these elements as a solid with a defined and uniform thickness (**Fig. 4D, S10E**). The two physics interfaces were coupled by having them solve for the same dependent displacement variables. All components were modeled as simple linear elastic materials with linear behavior that follows Hooke’s law.

### Geometry of the FeCO model

The geometry of the model was based on the x-ray reconstruction and confocal imaging of FeCO neurons, tendons, and other structures. The medial tendon was modelled as a 250 µm cable with a circular cross section with a radius of 2 µm (**Fig. 4D**). Each fibril emanating from the tendon connecting to a cap cell was modelled as a cable with a circular cross section with a radius 0.2 um. Each dendrite was modelled as a 50 um string with a thinner circular cross section with a radius of 0.2 µm. Both the tendon and fibrils were modelled as being constrained to remain straight, to capture their higher bending stiffness. 20 dendrites were modelled as laying parallel to each other and at an angle of 168° to the long axis of the medial tendon. 3 dendrites were also modelled as lying in parallel to the medial tendon to represent the claw cells at the most proximal end of the FeCO. In most models, we see relatively low bending in the truss structures as their shape is stabilized by surrounding tissue.

We used a single vertex at each end of dendrites to model the position of each cap and claw cell. We treated each cell as a point mass. The mass of each cell was set by assuming that they are spheres with a density of 1200 kg/m^3^ and radius of 4 µm for the claw cell and 2 µm for the cap cell. Fibrils were modelled by attaching the proximal tip of the medial tendon to each cap cell. In this geometry, fibrils joining the tip of the medial tendon to the cap cells form a radiating array with an angular gradient from 168° to 180° similar to that observed in the real FeCO.

Finally, we embedded this truss structure in a solid with a uniform thickness of 10 µm, length of 660 µm, and width of 80 µm. The FeCO is surrounded by a clear and low density tissue, resembling extra-cellular matrix, within the femur. The geometry of this solid roughly approximates the shape of the tissue around the FeCO. The edges of the solid are chamfered to retain the rounded structure of the leg and to prevent accumulation of stresses in corners. The solid is also narrowed proximally near the FeCO, where the femur connects to the next leg segment. The truss modelling the tendon is placed so that it is symmetrically embedded within the height of the tissue, and the entire medial tendon is always within the tissue space.

The model was meshed using 20 edge elements for each truss, i.e. for the medial tendon, each fibril and dendrite. The solid was meshed using triangular elements using COMSOL’s physics-controlled meshing at a finer mesh size. This uses a minimum element of 0.08 µm and allows a maximum element size of 24.4 µm. The model uses smaller elements near the FeCO, and in the narrower corners of the solid, to more carefully calculate strains and stresses as they develop in these regions. The complete geometry was represented by ∼6717 triangle elements and 1589 edge elements.

### Material properties of the FeCO model

We treated all parts of the model as linear elastic materials. Most insect cuticle lies within a narrow range of densities of ∼1200 kg/m^3^ (Vincent and Wegst, 2004) and all parts were modelled with this density. Most materials in nature have a Poisson’s ratio of 0.3, which was similarly uniformly applied. The exact Young’s modulus of the medial tendon is not known. Like the arculum, UV fluorescence data suggests that it contains some resilin. Again, since this is likely to be the stiffest material in this tissue, and therefore dominate its stiffness value, we used 1.8 MPa as the Young’s modulus for the medial tendon and for the thinner fibrils arising from it (Frantsevich et al., 2019; Gosline et al., 2002). For the ciliated dendrites of the claw cells, we used 178 kPa, an estimate of ciliary Young’s modulus made by previous authors (Rydholm et al., 2010). We also tested the effect of changing the stiffness of these elements (**Fig. S10F**).

We do not know the exact Young’s modulus of the tissue surrounding the FeCO. However, Young’s modulus of soft biological tissues are known to range from ∼100 Pa (mammalian brain) to several MPa (mammalian cartilage) (Levental et al., 2007). We expect the internal tissue of fly legs to be relatively soft. We tested the behavior of our model when the solid around the FeCO was set to a range of Young’s moduli between 100 Pa and 2 kPa. We found that when the modulus was set to 1 kPa, it reasonably reproduced the motions of cap and claw cells observed experimentally (**Fig. 4F, S7B, S8B, D**). Thus, we used this value in all models. This supports the idea that the internal tissue in the fly leg has to be relatively soft compared to the tendon and the dendrites for this system to function as observed.

We found that tissue stiffness interacts with the modelled thickness of the solid. If the solid’s thickness is increased by an order of magnitude (to 100 µm), then the Young’s modulus of the solid needs to be decreased by an order of magnitude (100 Pa) to recover the same level of cell displacement and strain gradient seen in the thinner stiffer solid (10 µm, 1 kPa). This contrasts with the behavior near the immobile tissue wall (**Fig. S10E**). This is because in this 3^rd^ dimension in the 2D model, there is no mechanism by which to set the wall as being immobile, and no drag like effects from the wall are observed. What we observed is that in conditions where the tissue edge is mobile, thinner tissues will allow greater motion, and thicker tissues resist motion. In the real tissue, the stiffness of the extracellular matrix is likely tuned by inclusion of different concentrations of structural protein (Levental et al., 2007). We expect that in the real system, the tissue’s width, thickness, and stiffness are appropriately matched.

We applied a high level of material damping to both the truss and solid material (isotropic loss factor of 0.8) to account for the high viscous damping encountered by small structures immersed in fluid. For the sake of simplicity, we ignored any material anisotropy in this tissue. In the real system, given the orientation of the fibers within the matrix, and the neurons juxtaposing the claw cells, there is likely some anisotropy. However, since these properties are difficult to estimate, we begin with a simplified system where soft isotropic tissue surrounds the FeCO and found that it can still reproduce most of the behavior of interest. Further tuning of the model might recover all measured features of the mechanics, but at the cost of further assumptions, which would lower the explanatory power.

### Boundary conditions of the FeCO model

We embedded and coupled the truss structure to the solid in the equilibrium position but otherwise left it unconstrained. We expect the tissue surrounding the FeCO to be bounded by other stiffer tissue along the edge and likely has a no-slip condition, i.e. is fixed to the boundary and does not move across it easily. Therefore, the solid that modelled the tissue surrounding the FeCO had fixed external boundaries, i.e. the solid could not move at the edges.

### Application of forces to the FeCO model

To simulate the forces applied to the claw cells in the FeCO by the slow flexion and extension of the tibia, we measured the movement of the center of mass of the arculum from flexion to extension from 12° to 176° at the rate of 3.28 °/s over 50s (**Fig. 3G**). We also measured the angle of the arculum over the same movement (**Fig. S6B**). From this, we can calculate the position of the attachment of the medial tendon to the arculum. Thus, we know the position of the distal end of the medial tendon as the leg joint is flexed/extended.

To identify a tibia position in which there are no net forces on the cap/claw cells in the FeCO, we held the tibia at different angles and relieved tension on the FeCO by cutting the medial tendon. We found that the cap cells moved in the proximal direction at all tibia angles, suggesting that FeCO is stretched distally even when the leg is fully extended (**Fig. S9B, C**). As expected from the movement of the medial tendon, the movement of the cells after the tendon cutting was larger when the tibia was flexed. Similar to the movements of the cap and claw cells during tibia flexion (**Fig. 4E**), at each tibia angle, the distal cells moved more than the proximal cells when we cut the tendon (**Fig. S9B, C**). Together these observations suggest that the FeCO cells are held under tension by being pulled upon by the medial tendon.

We used the information from the tendon cutting experiment and applied tension to the model FeCO by displacing the distal end of the medial tendon towards the distal direction. We chose the distance to move the medial tendon based on the cell movements during the cutting of the medial tendon with tibia held in the flexed position (**Fig. S9B**). In these experiments, the most distal cap cell moved a maximum of 35 um to its equilibrium position. Therefore, we moved the medial tendon by a distance that moved the most distal cap cell in our model an equivalent distance (35 um), replicating the tension in the flexed position. Next, we moved the distal end of the medial tendon in a way that resembled the observed movement of this point on the arculum as the leg moved from flexion to extension. At full extension, we found that the most distal cap cell was 9 um from the equilibrium position which was similar to the observations from the cutting experiment (**Fig. S9B**), validating our model further.

The application of the forces to the model is illustrated in **Fig. S10B**. The model was initiated at a hypothetical neutral position (from 0-3.5s), then tensioned (from 3.5-6.5 s), i.e. the distal end of the medial tendon was drawn back. The model was allowed to equilibrate into this tensioned position (6.5-10 s), since moving the FeCO from its equilibrium position will generate a large transient strain in the system, which is not relevant to our question. After the transient strain had decayed and the full steady state strain had formed, we started displacing the medial tendon in the model. It was displaced with the same dynamics as were observed in the real data, thus replicating the observed forcing regime in the model. In all subsequent analyses, we considered steady-state behavior of the model.

### Development of a strain gradient in the FeCO model

We found that the movement of cap and claw cells that we observe can be captured if the model has three important features, 1) a fan-like array of tendon fibrils connecting the medial tendon to the claw cell dendrites (**Fig. S10C, D**), 2) tissue that is relatively soft (**Fig. S10E**), and 3) tendon elements and dendrites that are stiffer than the surrounding tissue (**Fig. S10F**). The resulting relative motion of the cap and claw cells lead to a strain gradient that is highest in the distal most dendrite and then decays along the FeCO array (**Fig. 4G, H, S10D, F**). Additionally, we find that dendritic strain peaks when the tibia is flexed and reduces from this level as the tibia is extended (**Fig, 4G, H, S10D, F**).

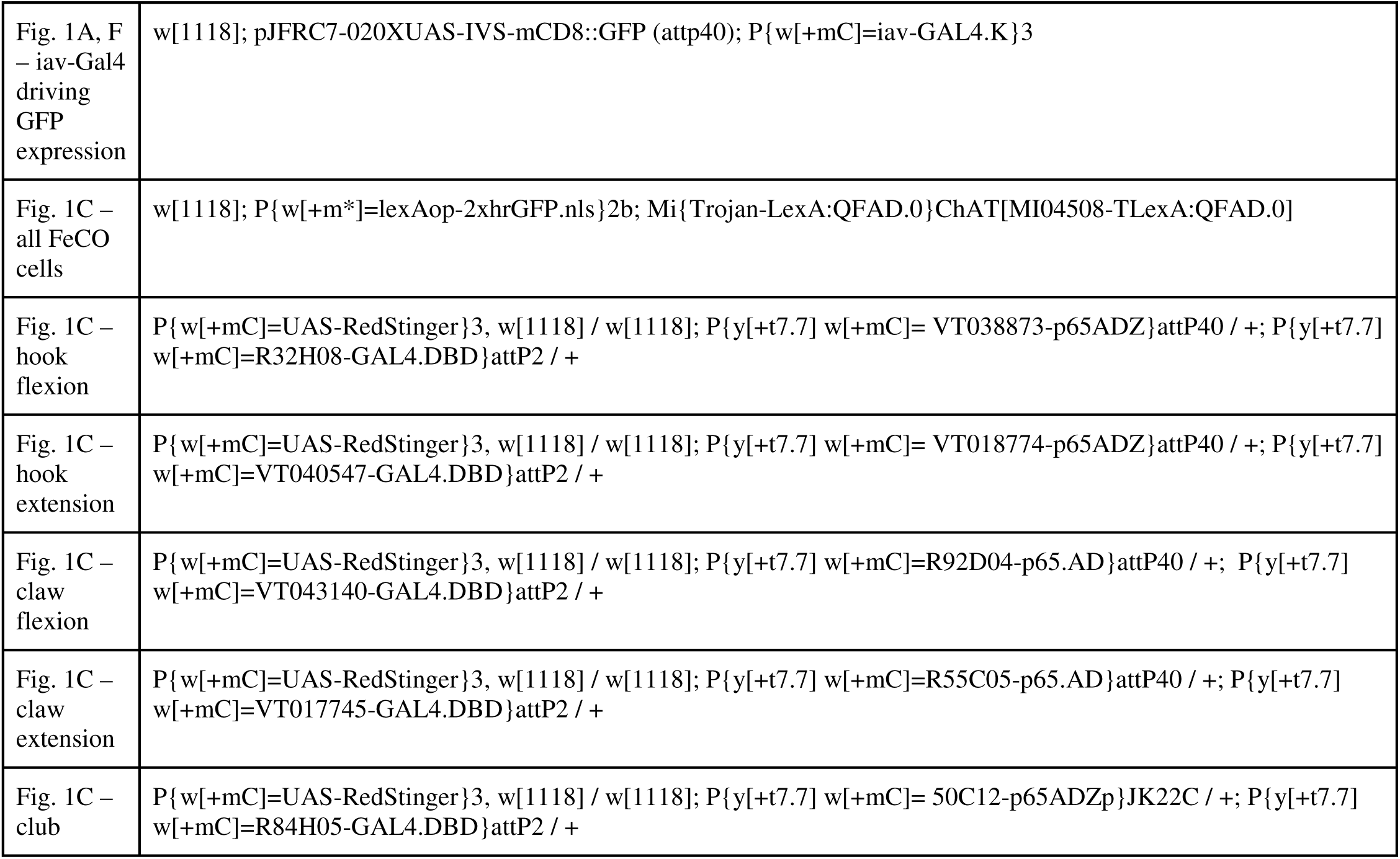
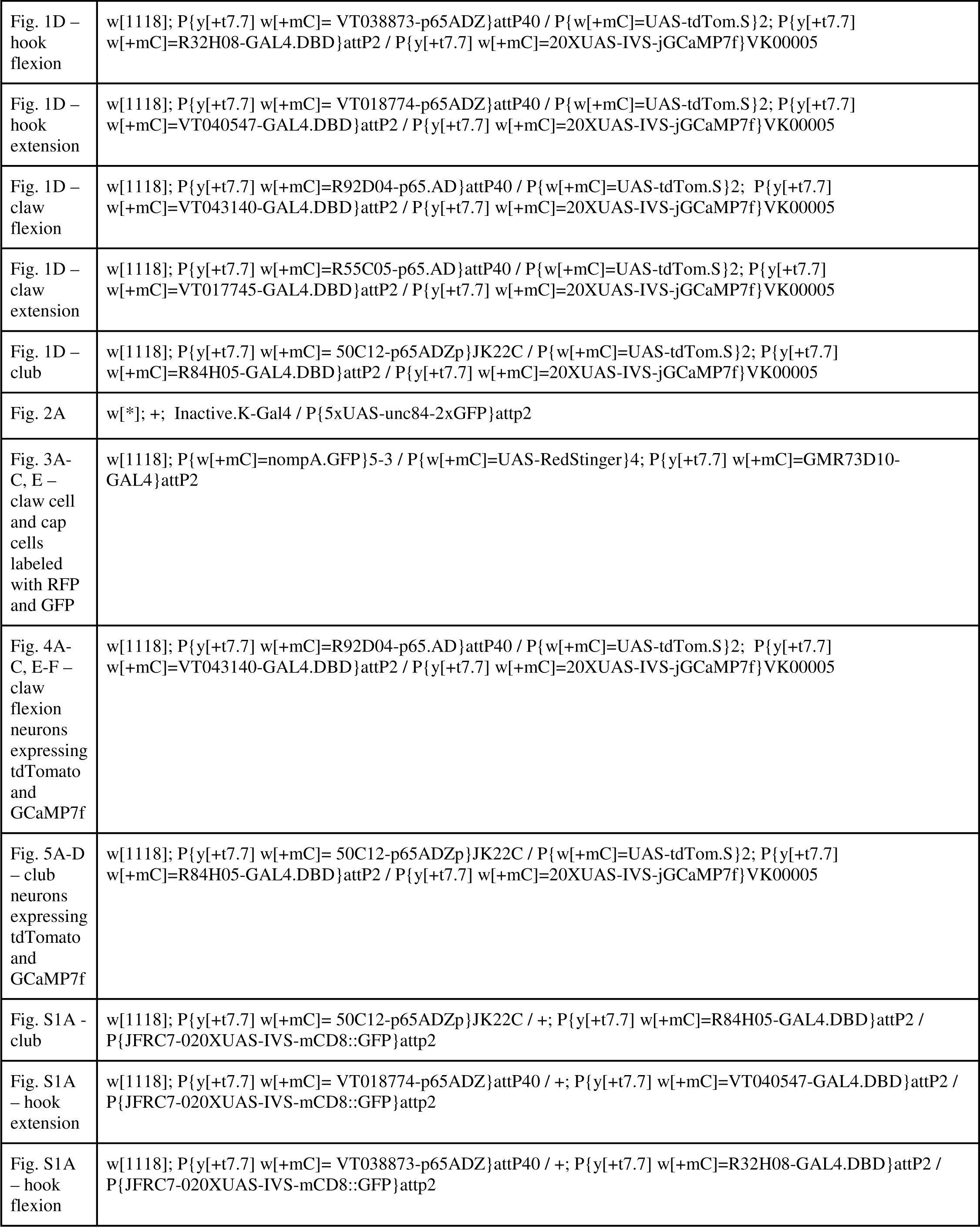
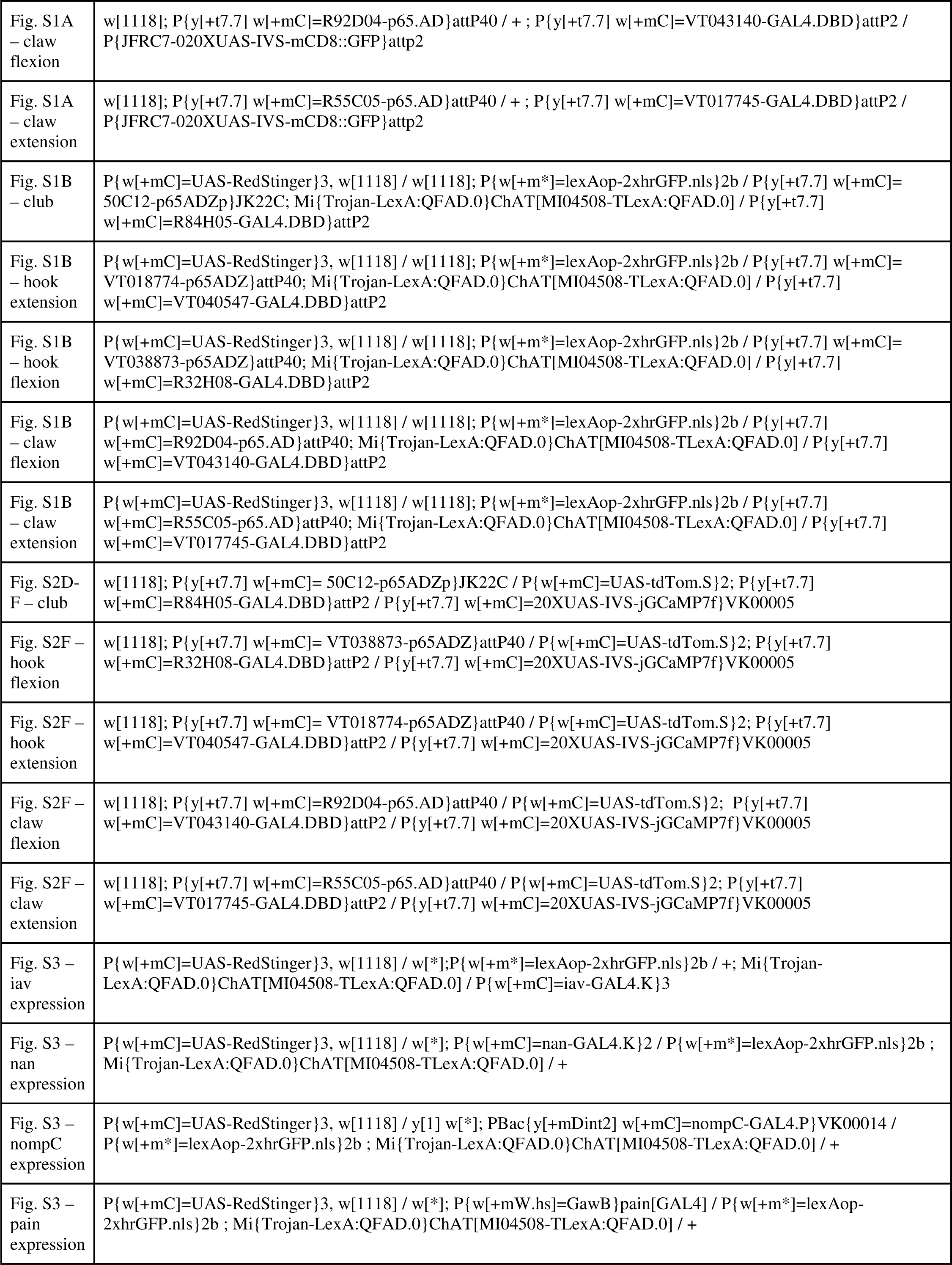
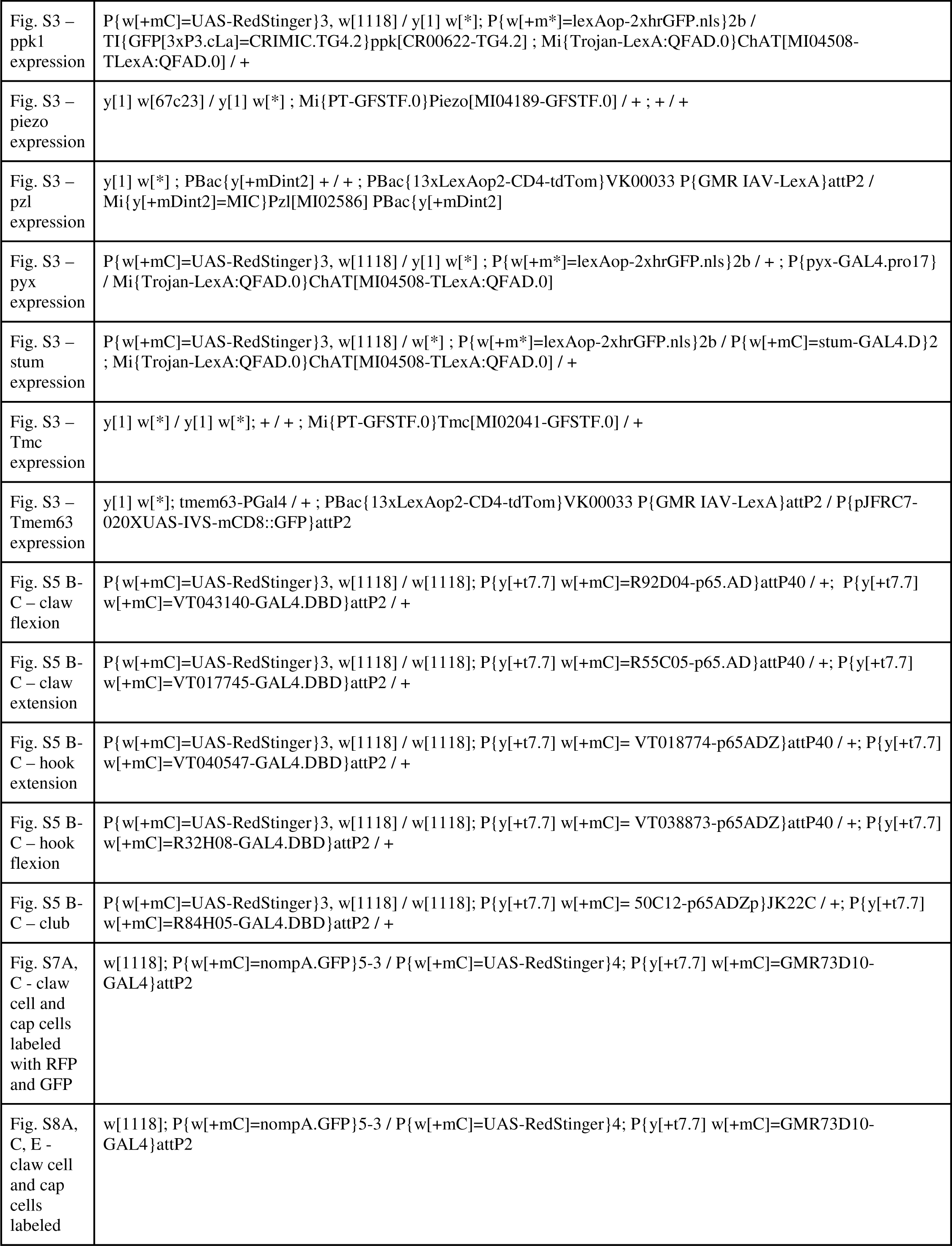
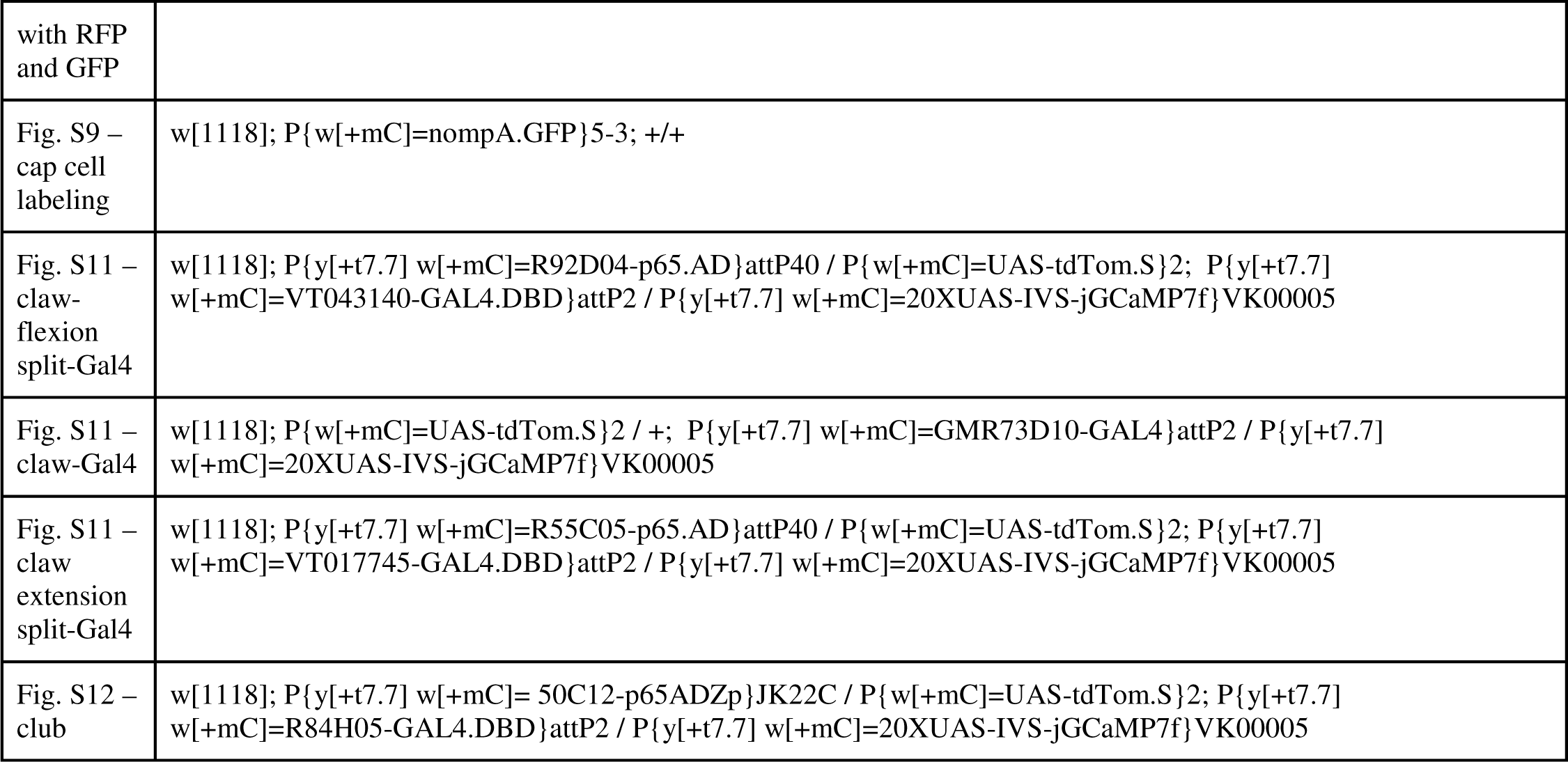
Table of genotypes

